# Translatome profiling reveals opposing alterations in inhibitory and excitatory neurons of Fragile X mice and identifies EPAC2 as a therapeutic target

**DOI:** 10.1101/2025.04.21.649817

**Authors:** Anand Suresh, Nazim Kourdougli, Jessie E. Buth, Carlos A. Sánchez-León, Lauren T. Wall, Anne T. Tran, Soledad Miranda-Rottmann, Roberto Araya, Michael J. Gandal, Carlos Portera-Cailliau

## Abstract

Symptoms of Fragile X Syndrome (FXS), the leading monogenic cause of intellectual disability and autism, are thought to arise from an excitation/inhibition (E/I) imbalance. Here, we leverage cell type specific mRNA sequencing to profile molecular alterations of cortical excitatory and inhibitory neurons in *Fmr1* knockout (KO) mice, integrating transcriptomic results with circuit and behavioral readouts to prioritize novel therapeutic targets. Differentially expressed genes (DEG) were largely upregulated in *Camk2a* expressing excitatory neurons but downregulated in *Pvalb*-expressing inhibitory neurons, and the underlying signaling pathways were often altered in opposite directions. Among the 184 DEGs that were concordantly dysregulated across both cell types, only *Rapgef4* (Epac2) was also an FMRP target, an ASD risk gene and brain enriched. EPAC2 has been implicated in synaptic maturation and plasticity. Systemic administration of an EPAC2 antagonist restored cortical circuit function in *Fmr1* KO mice and ameliorated sensory behavioral phenotypes. EPAC2 is a potential target for therapy in FXS.

## INTRODUCTION

Fragile X Syndrome (FXS) is a prototypical neurodevelopmental condition (NDC) characterized by intellectual disability, attention deficit, social anxiety, atypical sensory processing, and autistic traits^1^. It is the most common single gene cause of intellectual disability and autism spectrum disorder (ASD), affecting 1 in 2,000 children^2^. FXS is caused by mutations in the *FMR1* gene, typically a CGG triplet repeat expansion within the 5’ untranslated region of the promoter region^3,4^. This results in transcriptional silencing of the gene and the near complete loss of Fragile X Messenger Ribonucleoprotein (FMRP). FMRP is an RNA binding protein that is enriched in neurons and plays diverse functions in different cellular compartments (e.g., nucleus, dendrites, axons), including regulation of mRNA translation at the synapse^5^.

Since the discovery of the genetic basis of FXS in 1991, several clinical trials have been unsuccessful and, presently, there exist no specific treatments for FXS^6,7^. Thus, there is an urgent need to rethink therapeutic strategies for FXS. One potential avenue is to leverage RNA sequencing (RNA-seq) to systematically characterize changes in gene expression, providing potential insights into molecular pathophysiology that might reveal novel candidates for targeted therapy. Studies of the translatome (RNA transcripts bound to ribosomes that are actively being translated) could be uniquely useful in FXS because FMRP is an mRNA binding protein that regulates the translation of ∼1,200 mRNAs, including those encoding for proteins important for neuronal development and synaptic function^8,9^, a large fraction of which are linked to ASD and other NDCs, underscoring the importance of FMRP in brain function^10–13^.

Although several studies have characterized changes in the translatome after loss of FMRP in mice, they focused on excitatory neurons of the hippocampus. To date, no study has directly compared changes in excitatory vs. inhibitory neurons or across cortical regions^14–18^. Our previous RNA-seq study probed only GABAergic inhibitory neurons from the medial ganglionic eminence (MGE)^19^. A better understanding of how FMRP loss affects the translatome in these cell types is critical within the general framework of the E/I imbalance theory of FXS and other NDCs^20,21^, because there exists some debate as to which cell type, excitatory or inhibitory, is primarily affected in FXS^22,23^. Although several studies have reported a lower density and/or hypoactivity of parvalbumin (PV) interneurons in *Fmr1* KO mice^19,24–26^, others have argued that this could represent a secondary, compensatory response^27^. Identifying which differentially expressed genes (DEGs) are unique or shared between these two cell types would provide important insights into cell type-specific circuit dysfunction and help discover potential molecular targets for therapy in FXS.

We used an RNAseq approach with RiboTag^28^ to isolate the translatome from *Pvalb*-expressing interneurons and *Camk2a*-expressing excitatory neurons from both WT and *Fmr1* KO mice. We also compared the translatome of primary somatosensory (S1) and primary visual cortex (V1) to test for regional differences or convergence. We find hundreds of DEGs in excitatory neurons but 5-fold fewer in PV interneurons. Although regional differences were negligible, differences across cell types were profound. Consensus network analysis revealed an excitatory neuron module exhibiting convergent enrichment for both previously defined direct FMRP mRNA targets^10,29^ and ASD risk genes^30,31^, characterized by chromatin and synaptic gene pathways. Intriguingly, only 193 DEGs were shared between excitatory and PV neurons (∼6% and ∼36% of total DEGs, respectively), and among these only one gene, *Rapgef4* (a.k.a., *Epac2*), was also an ASD risk gene, a target of FMRP, and highly expressed in the brain. EPAC2, a cAMP dependent guanine-exchange factor, is a known regulator of synapse turnover and stability^32,33^. We found that chronic treatment of *Fmr1* KO mice with ESI-05, a specific EPAC2 inhibitor, improved circuit and behavioral phenotypes related to sensory processing.

## RESULTS

### Profound differences in translatome dysregulation between excitatory and inhibitory neurons of *Fmr1* KO mice

We used RiboTag RNA-seq^28^ to deeply profile the cell type and brain region-specific translatomes of excitatory and PV inhibitory neurons between *Fmr1* KO mice and WT controls. We generated triple transgenic *Camk2a*-*Cre*^+/−^; *Rpl22*^HA/HA^ and *Pvalb*-*Cre*^+/−^;*Rpl22*^HA/HA^ mice of both genotypes at 10-14 weeks, collected region-specific tissue (S1, V1), and isolated ribosome-bound RNA (**Figure 1A**; see Methods). We collected samples from mice of both sexes (35 mice: 15 *Camk2a*-Cre, 20 *Pvalb*-Cre; 18 WT, 17 *Fmr1* KO) and sequenced at a depth of 50 million reads per sample. Principal component analysis and hierarchical clustering of gene expression identified clear separation of samples according to both cell type and brain region (**Figure 1B and Figure S1A-B**). The expression of known markers for excitatory (*Camk2a*, *Slc17a7*) and PV neurons (*Pvalb*, *Gad2*), as well as markers of other interneuron subtypes (*Sst* and *Vip*), confirmed the cell type-specificity of this approach (**Figure 1C**). As expected, *Fmr1* expression was substantially reduced across all samples from *Fmr1* KO mice relative to WT controls (**Figure 1D**). We also cross-referenced with cell type specific marker genes from the Allen Brain single-cell dataset^34^. The *Camk2a* and *Pvalb* translatomes exhibited strong and selective enrichment for their respective cell markers both in comparison to the bulk transcriptome and to other cell types (**Figure S1A-B**). Finally, using immunohistochemistry we confirmed that hemagglutinin (HA) expression (for ribosomal *Rpl22^HA/HA^*) was restricted to the target cell type (**Figure S1D**).

**Figure 1.**
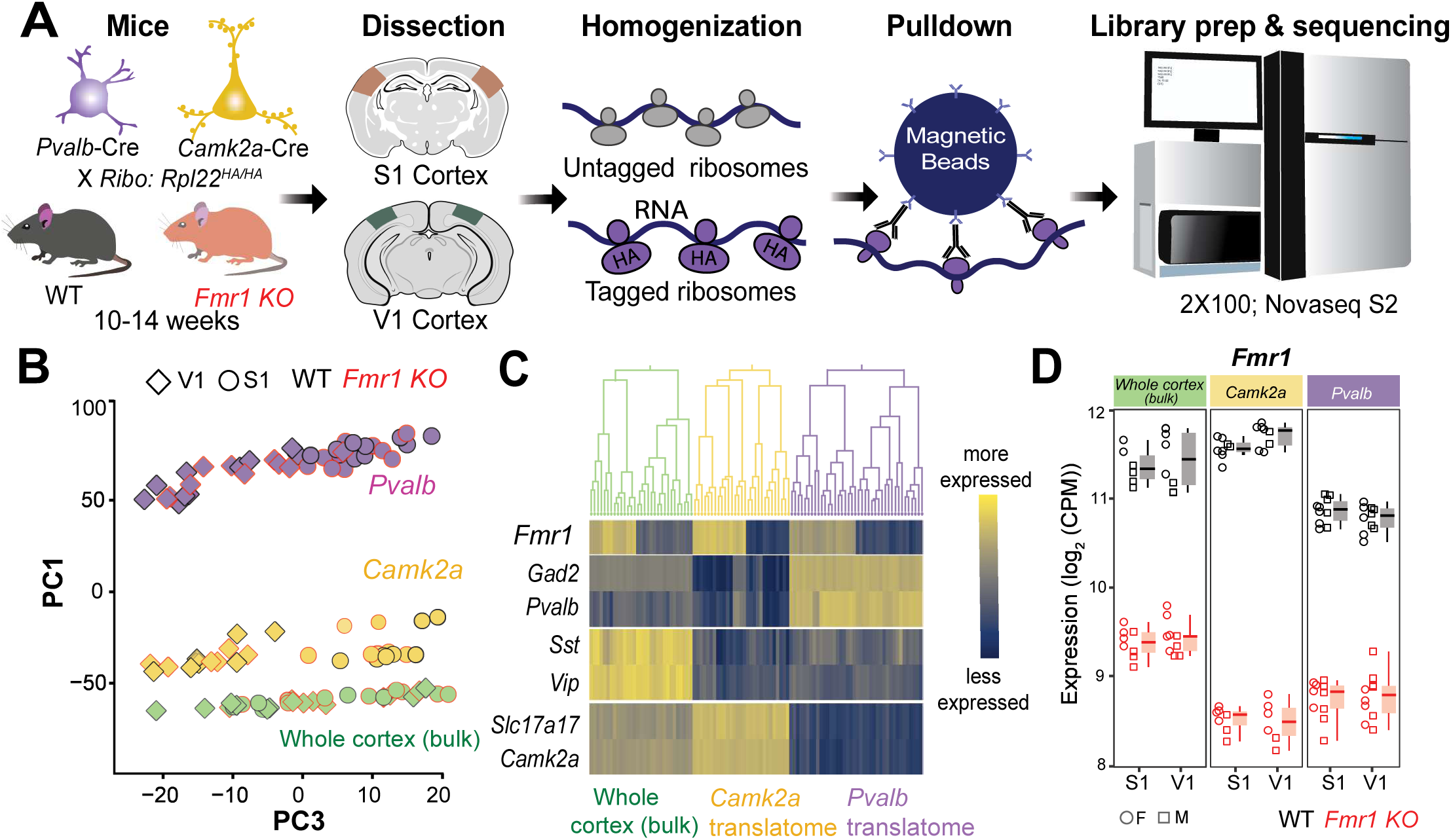
RiboTag approach to obtain cell type-specific translatome from excitatory and PV neurons using *Camk2a*-Cre and *Pvalb*-Cre *Fmr1* KO and WT mice. **(a)** Cartoon of Ribotag RNA-seq approach in *Camk2a*-*Cre* and *Pvalb-Cre* mice. **(b)** Principal component (PC) analysis of normalized gene expression shows clear separation of samples by cell type and region (PC2, not shown, also separated samples by cortical region). **(c)** Excitatory and PV neuron marker genes are enriched in the *Camk2a* and *Pvalb*-specific translatomes, respectively, but markers of other interneuron subtypes (*Sst*, *Vip*) are not. **(d)** *Fmr1* expression is substantially reduced in *Fmr1* KO samples. S1 whole cortex: adjusted p; S1 whole cortex = 7.45e-19, V1 whole cortex = 6.35e-15; S1 *Camk2a* = 4.25e-38; V1 *Camk2a* =6.87e-33; S1 *Pvalb*: = 5.59e-30; V1 *Pvalb* =1.32e-28. Data represented as median±1.5*IQR.

We next quantified patterns of differential gene expression following loss of FMRP in *Camk2a*-Cre^+/−^; *Rpl22*^HA/HA^ mice (8 WT, 7 *Fmr1* KO) and *Pvalb*-Cre^+/−^;*Rpl22*^HA/HA^ mice (10 WT, 10 *Fmr1* KO) using *limma voom*^35^, across both cortical areas (FDR=0.05). In excitatory neurons (*Camk2a* translatome), we identified 3,046 DEGs (1,531 up-regulated and 1,515 down-regulated) in *Fmr1* KO mice (**Figure 2A-B**), with *Fmr1* as the top downregulated gene. Among the top DEGs in excitatory neurons were several that had been identified in previous RNA-seq studies, including *Atmin*^14,17,19^, *Setd2*^36^, *Ank3*^37^, and *Wdr81* (up), which is highly expressed in the brain and associated with an autosomal recessive form of intellectual disability and ataxia^38^. Notably, *Rptor* (Regulatory-associated protein of mTOR complex 1) was also upregulated, consistent with increased protein synthesis and enhanced mTOR activity in FXS^39,40^.

**Figure 2.**
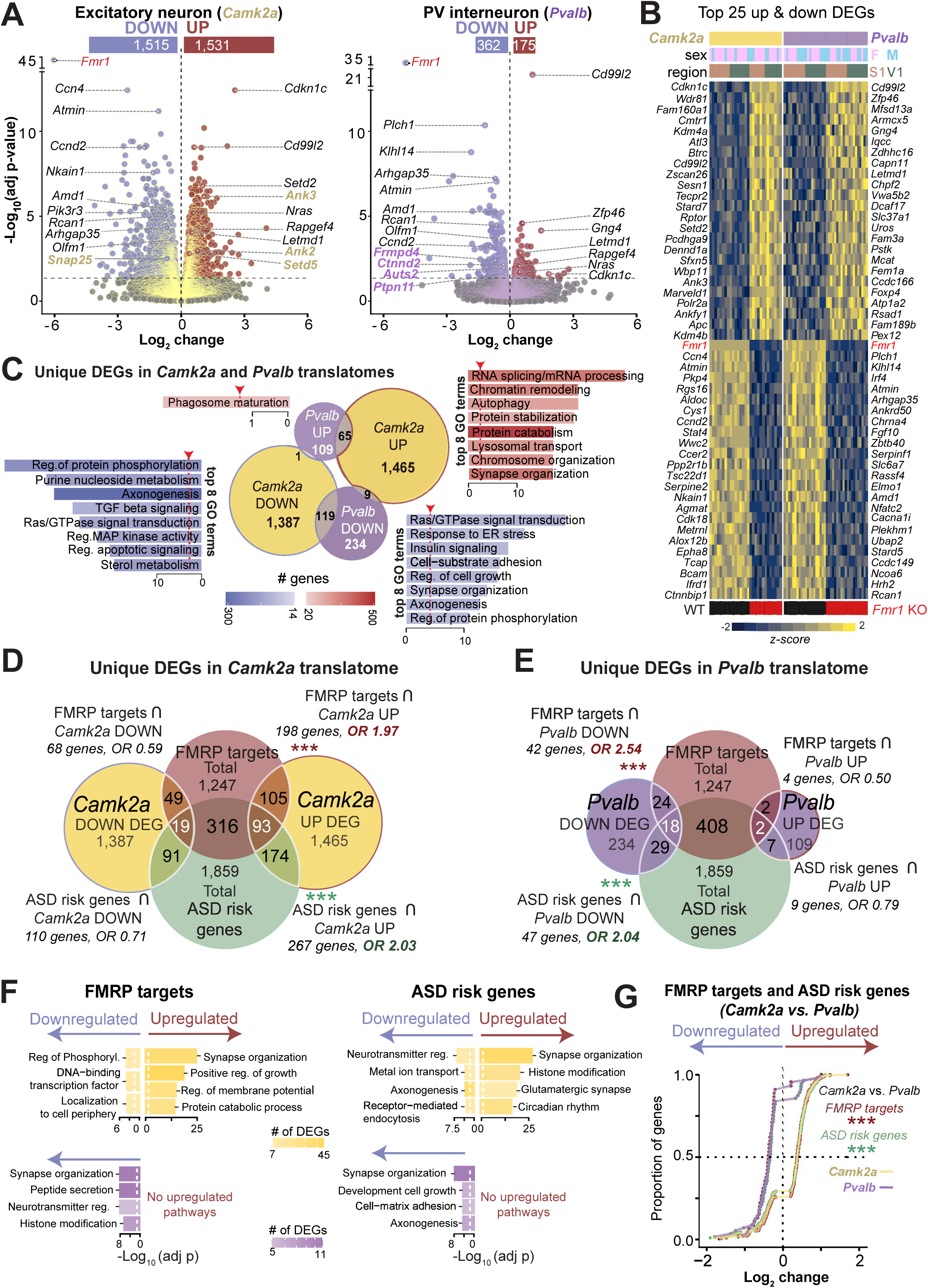
Unique changes in differential gene expression in translatome of excitatory vs. PV inhibitory neurons of *Fmr1* KO mice. **(a)** Volcano plot of DEGs (*Fmr1* KO relative to WT) in the *Camk2a* excitatory translatome (13 *Fmr1* KO, 16 WT mice) and PV inhibitory translatome (20 *Fmr1* KO, 20 WT mice). Upregulated genes are indicated in red and downregulated genes in blue. Horizontal grey dashed line represents the negative log of adj.p (FDR=0.05). **(b)** Expression of top twenty-five up and down DEGs sorted by statistical significance. Heatmaps represent changes of expression relative to WT mice (expressed as Z scores). **(c)** Comparison of uniquely up-or downregulated DEGs and the top 8 associated GO terms. The red dotted vertical lines and arrows represent the log of adjusted p<0.05. Note the significant and positive association for *Camk2a* Up∩*Pvalb* Up (Odd’s ratio, OR=6.43, p=2.16e-25) and *Camk2a* Down∩*Pvalb* Down (OR=5.56, p=6.3e-40). ***p<0.001, Fisher’s exact test for panels C-E. **(d)** Overlap between DEGs unique to *Camk2a* translatome and FMRP target genes or ASD risk genes. There was a significant positive association between upregulated DEGs in *Camk2a* neurons and both FMRP targets (p=1.23e-14) and ASD risk genes (p=1.46e-19) but not in downregulated DEGs (FMRP targets: OR=0.58; ASD risk genes: OR=0.71). We also confirmed the known strong association between FMRP targets and ASD risk genes (OR=8.06, p=1.34e-176). **(e)** Same as D but for *Pvalb* DEGs. Note the significant positive association between downregulated DEGs unique to *Pvalb* translatome and both FMRP target genes (p=1.24e-6) and ASD risk genes (p=5.26e-5), but not for upregulated DEGs. **(f)** GO enrichment for genes that overlapped between FMRP targets or ASD risk genes and *Camk2a*-unique DEGs (top, yellow) or downregulated *Pvalb*-unique DEGs (bottom, purple). The white dotted vertical lines represent the log_10_ adjusted p<0.05 (FDR=0.05). No pathways for upregulated genes in *Pvalb*. **(g)** Cumulative distribution of *Camk2a* DEGs (yellow lines) and *Pvalb* DEGs (purple lines) that overlapped with FMRP targetd (green outline) or ASD risk genes (red outline). Horizontal dotted lines indicate the median (0.5) and vertical dotted lines indicate log fold of 0 (*Camk2a* vs. *Pvalb* FMRP targets, p<2.3e-13, *Camk2a* vs *Pvalb,* ASD risk genes p<2.3e-13 Wilcoxon signed-rank test). ***p<0.001

In PV neurons (*Pvalb* translatome) we identified only 537 DEGs, far fewer than in *Camk2a* translatome, with two-fold more downregulated (362) than upregulated genes (175). Apart from *Fmr1*, the top DEGs in PV interneurons included previously identified genes like *Atmin, Arghap35*^17^ and *Cacna1i,* which encodes for the pore forming alpha-1 subunit of the Cav3.3 voltage-dependent calcium channel and is associated with ASD^41^ and NDCs^42^.

We next sought to identify genes (and pathways) uniquely affected within each cell type (**Figure 2C**; see Methods). We found 2,852 uniquely dysregulated genes in *Camk2a* neurons (1,465 up and 1,387 down) and 343 uniquely dysregulated genes in *Pvalb* neurons (109 up and 234 down). *Camk2a*-specific upregulated DEGs were enriched for ‘RNA splicing/mRNA processing’, ‘Chromatin remodeling’ and ‘Autophagy’ gene ontology (GO) terms, whereas ‘Regulation of protein phosphorylation’, ‘Purine nucleoside metabolism’ and ‘Axonogenesis’ were the top enriched pathways among downregulated genes (**Figure 2C**; see full list in **Figure S2A-B**). For *Pvalb*-specific DEGs, the top GO terms for downregulated genes were ‘Ras/GTPase signal transduction’, Response to ER stress’, and ‘Insulin signaling’, while only Phagosome maturation’ was identified for upregulated genes (**Figure 2C and Figure S2C-D**).

These initial findings reveal that loss of FMRP exerts distinctly contrasting effects on the translatomes of excitatory and inhibitory neurons. In PV neurons, there is an overall downregulation of gene expression, whereas in *Camk2a* neurons the dysregulation is more balanced. Overall, biological pathways are affected in different, many times opposite ways between the cell types (**Figure S2**). Notably, ‘Insulin signaling’ was upregulated in *Camk2a* neurons but downregulated in *Pvalb* neurons, which may reflect opposite energetic demands of each cell type in FXS. Together with the opposite effects on ‘Synapse organization’, ‘Chromatin remodeling’, and ‘Autophagy’ (**Figure S2**), the overall picture is consistent with the hypofunction of PV neurons in *Fmr1* KO mice that recent studies have reported^19,21,26,43^.

### DEGs are enriched for FMRP targets and ASD risk genes but often in opposing ways in each cell type

FMRP plays a crucial role in neuronal maturation and plasticity and is known to function as a translational repressor of ∼1,250 target transcripts^10,12^, many of which overlap substantially with known ASD risk genes^11,13,44^. Thus, we next characterized the overlap between cell type-specific DEGs and these known FMRP targets (1,247 genes), as well as with ASD risk genes (1,859 genes) obtained from the SFARI database and the Geisinger developmental brain disorder database. Among the *Camk2a*-specific DEGs, there was significant association of upregulated genes (but not downregulated genes) with both FMRP targets and ASD risk genes (odds ratio, OR=1.97 and 2.03, respectively; **Figure 2D**). In contrast, the opposite pattern was observed for *Pvalb*-specific DEGs, where only downregulated genes showed a significant association with FMRP targets and ASD risk genes (OR=2.54 and 2.08, respectively; **Figure 2E)**. Gene enrichment analysis for DEGs that were FMRP targets or ASD risk genes identified different pathways for *Camk2a*- vs. *Pvalb*-specific DEGs (**Figure 2F**). Intriguingly, synapse organization’ was affected in opposite ways in the two cell types (**Figure 2F**, **Figure S3A-B**). A SynGo analysis^45^ of these *Camk2a*-specific synaptic DEGs, identified ‘organization’ and ‘postsynaptic’ categories as the most prominently affected (**Figure S3C**).

When we compared fold-change differences for all DEGs in *Fmr1* KO mice, we found that in excitatory neurons most were upregulated, whereas in inhibitory neurons most were downregulated **(Figure S4A)**. This was also the case for DEGs that were also FMRP targets and ASD risk genes, and the differences were even more pronounced (**Figure 2G**, **Figure S4B-C**). This same pattern was also apparent when we plotted the functional enrichment using gene set enrichment analysis (GSEA; **Figure S4D**). Overall, this further suggests that loss of FMRP affects excitatory and PV inhibitory neurons in very different ways, often in opposite directions.

### Co-expression network analysis identifies cell type specific cellular networks in excitatory and inhibitory neurons

To more finely dissect the cell type specific biological networks dysregulated in *Fmr1* KO mice, we next conducted unsupervised gene co-expression network analysis using the weighted gene correlation network analysis (WGCNA) framework^46^. We constructed consensus networks across all samples, capturing shared variations within each cell type, which identified 29 distinct gene modules (**Figure 3A-B**). Of these, 10 were significantly dysregulated in excitatory neurons, but only 6 in PV neurons. Excitatory neurons showed larger effect size changes among co-expression networks than PV neurons, with only slight overlap in differentially expressed modules across cell-types (**Figure 3B**). More modules showed significant excitatory-specific (7) than PV-specific (2) dysregulation, once again highlighting how excitatory neurons are more affected by loss of FMRP.

**Figure 3.**
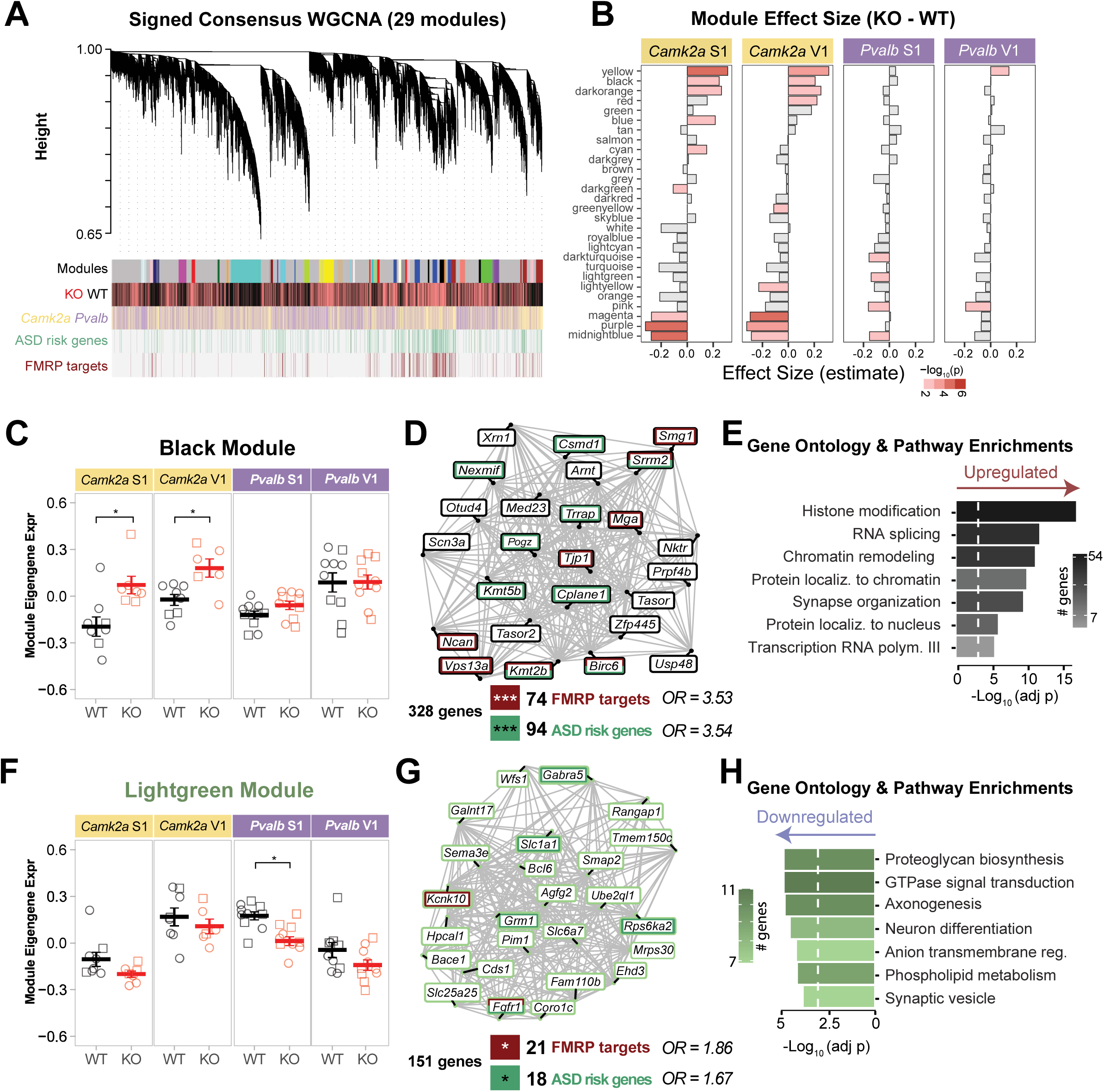
Consensus WGCNA analysis identifies cell type-specific modules enriched for FMRP targets and ASD risk genes. **(a)** Top: Gene dendrogram of consensus network (29 modules); Bottom: corresponding module colors, differential expression *Fmr1* KO vs. WT, relative expression in *Camk2a* vs. *Pvalb*. ASD risk genes, and FMRP target genes. **(b)** Effect size of genotype based on the eigengene expression for each module in different cell types and cortical areas. Bar size indicates the effect size (*Fmr1* KO vs. WT), while red hue intensity indicates significance, as −log_10_(adjusted p). **(c)** Black module eigengene expressions were significantly upregulated in a *Camk2a*-specific manner. (*Camk2a* S1 WT vs KO, p=0.011, V1 WT vs KO, p=0.012, *Pvalb* S1-V1 WT vs KO, p=0.9). **(d)** Top 25 hub genes for the black module showing FMRP targets (maroon) and ASD risk genes (green). The black module showed significant enrichment for FMRP targets (p=3.0e-20), and ASD risk genes (p=4.59e-24). Fisher’s exact test. **(e)** Gene ontology (GO) enrichment analysis of the black module (white dotted line marks the significance threshold). **(f)** Lightgreen module eigengene expressions were significantly downregulated in a *Pvalb*-specific manner and only in S1 (*Pvalb* S1 WT vs KO, p=0.011, V1 WT vs KO, p=0.9, *Camk2a* S1-V1 WT vs KO, p=0.9). **(g)** Same as *D* but for lightgreen module. These hub genes were significantly enriched for FMRP targets (p=0.014) and ASD risk genes (p=0.043). **(h)** GO enrichment analysis of the lightgreen module. *p<0.05, **p<0.01, ***p<0.001

Two modules were notable because they were significantly enriched for genes that were FMRP targets and ASD risk genes: the black module, which was specifically upregulated in *Camk2a* samples (FMRP targets: OR=3.53; ASD genes: OR=3.54; **Figure 3C-D**), and the lightgreen module (FMRP targets: OR=1.83; ASD genes: OR=1.68; **Figure 3F-G)**, which was specifically downregulated in *Pvalb* samples within S1.

Consistent with broader DEG analyses, black module genes were enriched for ‘histone modification, chromatin remodeling’, and ‘RNA splicing’ (OR=1.86, p=0.004), with several chromatin/histone modifiers and high-confidence ASD risk genes among top (hub) genes, including *Kmt2b*, *Kmt5b*, *Setd2* (**Figure 3E and Figure S5A**). ‘Synapse organization’ was another top upregulated pathway in the black module. Remarkably, genes involved in ‘chromatin remodeling’ were significantly overrepresented in excitatory neurons (OR=1.32, p=0.040), with a majority (60%) being upregulated (**Figure S5B-C**). Additionally, 40% (32/79) of these chromatin remodeling DEGs were ASD risk genes, FMRP targets, or both (**Figure S5C**).

In contrast, DEGs in the lightgreen module, which contained a smaller subset of FMRP target genes (21/151) and ASD risk genes (14/151), were associated with pathways related to neural development and function, such as ‘Axonogenesis’, Neuron differentiation’, ‘Synaptic vesicle’ and ‘Proteoglycan biosynthesis’ (**Figure 3G-H**). This is consistent with previous studies showing developmental delays of inhibitory neurons^21,22^, including impairments in the formation of perineuronal nets in FXS^25,47^.

Thus, WGCNA further underscores how loss of FMRP has a profound and preferential impact on biological gene networks of excitatory neurons, highlighting the clear cell type-specific differences in gene dysregulation in Fragile X mice.

### Minimal differences across cortical regions in the *Camk2a* and *Pvalb* translatomes

Various synaptic and circuit phenotypes have been observed in *Fmr1* KO mice in primary sensory cortices, including S1 and V1^48^. However, because these regions exhibit distinct cytoarchitecture and patterns of functional connectivity, are known to mature at different rates during development, and show distinct transcriptomic changes in ASD^49^, we assessed whether FMRP loss differentially affects the molecular signature of these two regions. We observed large differences between the translatomes of S1 and V1 in WT mice, with thousands of genes exhibiting regional patterns of differential expression across both *Camk2a* and *Pvalb* samples (**Figure S6A-B**). Yet, when comparing genotypes, we identified only two genes, *Smoc2* and *Fbxo33*, that were significantly dysregulated in a region-specific manner in excitatory neurons of *Fmr1* KO mice (**Figure 4A-B**), but none in PV interneurons (**Figure 4C**). This suggests that loss of FMRP affects S1 and V1 in a similar manner.

**Figure 4.**
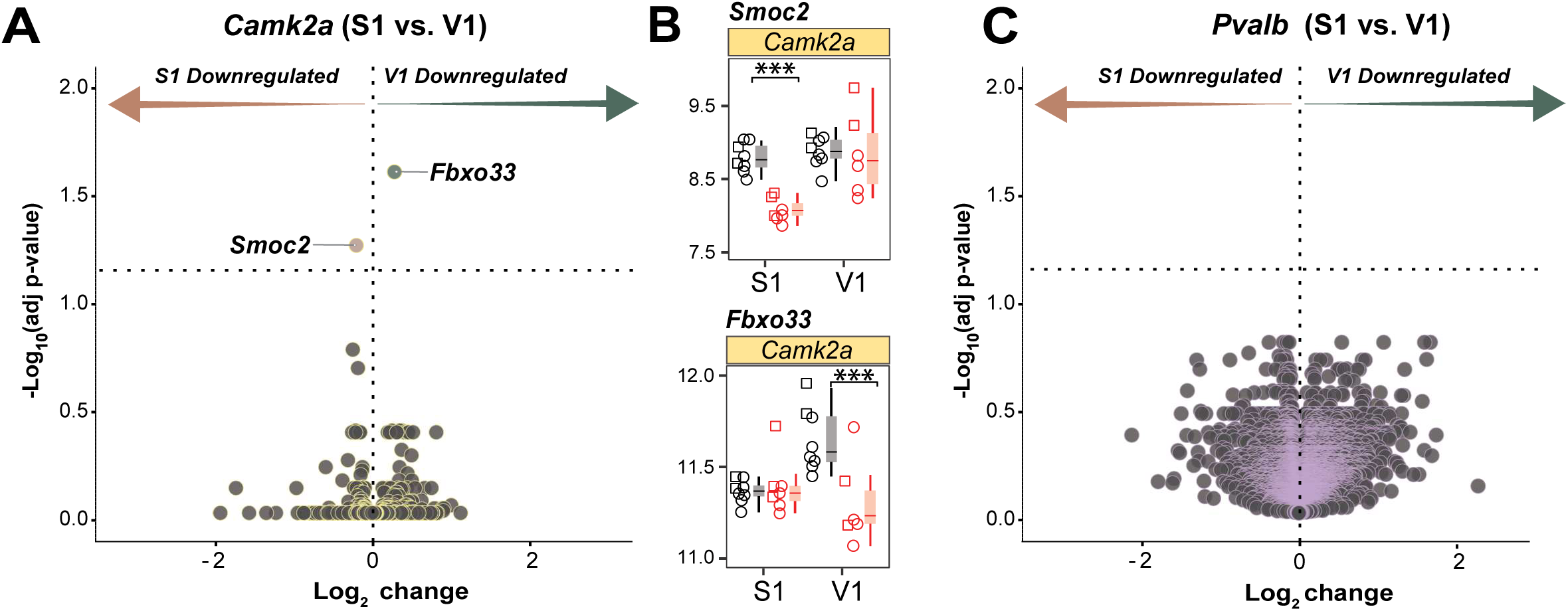
Negligible differences in differential gene expression across cortical areas. **(a)** Volcano plot shows DEGs in *Fmr1* KO mice that are specifically downregulated in S1 or V1 for the *Camk2a* excitatory translatome. *Smoc2* and *Fbxo33* are the only genes showing significant region-specific dysregulation (FDR=0.05). Horizontal dotted line marks the threshold of significance of p<0.05. **(b)** *Smoc2* and *Fbxo33* (from *A*) showed specific downregulation in *Fmr1* KO mice only in S1 or only in V1, respectively (adjusted p-value *Smoc2* S1=9.9e-08, *Smoc2* V1=0.7; *Fbxo33* S1=0.87; V1=1.39e-07). Data represented as median±1.5*IQR. ***p<0.001. **(c)** Same as *A* but for the *Pvalb* translatome. No genes show significant region-specific dysregulation.

### Comparison with a previous RNA-seq study of GABAergic interneurons reveals a developmental switch in the inhibitory translatome of *Fmr1* KO mice

We had previously characterized the translatome of interneuron precursors from the MGE (using Ribotag in *Nkx2.1*-Cre mice) and identified 2,089 DEGs in S1 of 2-week-old *Fmr1* KO mice^19^. Therefore, it was surprising that the number of DEGs in the *Pvalb* translatome of the present study in adult mice was much lower (537 DEGs). Still, we found that 95 DEGs in *Pvalb* samples (S1 only) overlapped above chance levels with the MGE translatome (OR=1.46), as did 468 genes in *Camk2a* translatome (OR=1.36; **Figure 5A**). Gene enrichment analysis identified no pathways for DEGs shared with the *Pvalb* translatome. In contrast, several up- and downregulated pathways were identified for those shared with the *Camk2a* (**Figure 5B**), including ‘Protein catabolism’, Autophagy’, ‘Histone modification’ and ‘Insulin signaling’ (**Figure 5B**), similar to what we had observed for *Camk2a*-unique DEGs. This was also evident in the distribution of DEGs according to fold-change, where the MGE translatome overlapped better with the *Camk2a* translatome than with the *Pvalb* translatome (**Figure 5C**). Tracking the trajectory of the *Nkx2.1* DEGs and PV DEGs across development from P15 to adulthood revealed that very few retained their directionality of dysregulation, while most ceased to be dysregulated (**Figure 5D-E**). Notably, several FMRP and ASD risk genes shifted from being upregulated in the *Nkx2.1* translatome to being downregulated in the PV translatome (**Figure S7A**). Despite these developmental shifts, there was a substantial positive correlation for DEGs across both samples (**Figure 5F**). These findings suggest that *Fmr1* loss of function affects inhibitory neurons differently across development, and that the PV translatome transitions into a hypofunctional phenotype in adulthood.

**Figure 5.**
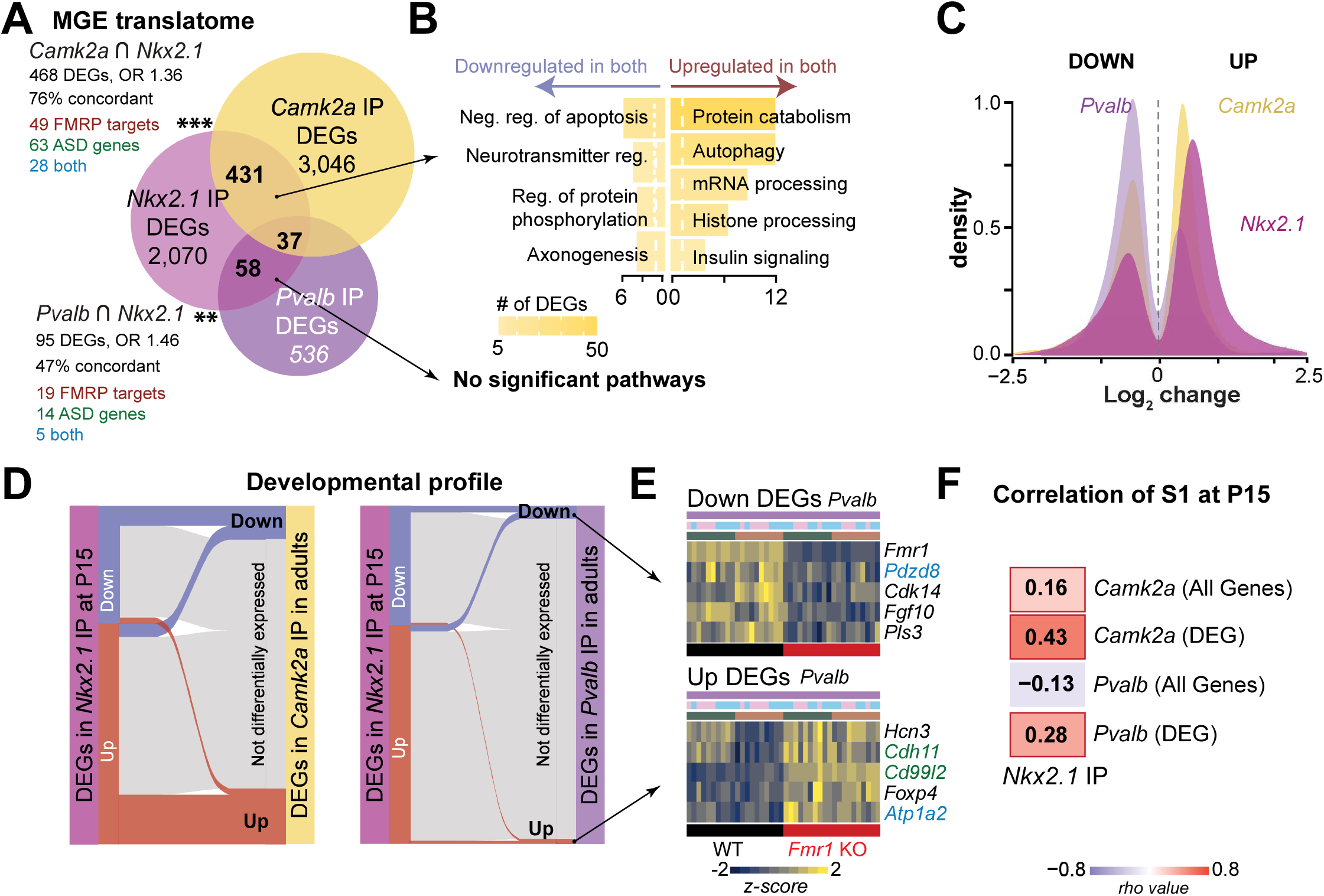
Comparisons of *Camk2a* and *Pvalb* DEGs with published *Fmr1* KO mouse MGE translatome and cortical proteomes, and with transcriptomes from human-derived FXS tissue. **(a)** Significant and positive association between DEGs (*Fmr1* KO vs. WT) from adult *Camk2a* and *Pvalb* translatomes and DEGs from P15 *Nkx.2.1* (MGE interneurons; Kourdougli et al., 2023) (*Camk2a*∩*Nkx2.1*: p=3.53e-07, *Pvalb*∩*Nkx2.1*: p=0.002). Fisher’s exact test for A, G, J-K. **(b)** GO enrichment of biological processes for shared DEGs from *A*. **(c)** Frequency distribution of fold change for DEGs in *Nkx2.1, Pvalb* and *Camk2a* translatomes. **(d)** Sankey plots tracking *Nkx2.1* DEGs and their subsequent fate in adult *Camk2a* (left) and *Pvalb* translatomes (right). **(e)** Top 5 up and down DEGs that retained directionality of dysregulation between *Nkx2.1* neurons at P15 and *Pvalb* neurons in adulthood. **(f)** Correlation coefficients between *Nkx2.1* translatome and either *Camk2a* or *Pvalb* translatomes (all genes or only DEGs). *p < 0.05, **p < 0.01, ***p < 0.001

### Significant overlap of *Camk2a* and *Pvalb* DEGs with previous proteomic studies of *Fmr1* KO mice and with RNA-seq studies of human FXS samples

We also compared our results with previous proteomic studies of cortical or hippocampal neurons^16,50–53^ (**Figure S7B**). We found that 201 DEGs from the *Camk2a* dataset and 30 DEGs from the *Pvalb* dataset (excluding *Fmr1*) were also dysregulated at the protein level (**Figure S7C**). Even though these overlaps did not reach significance (unsurprising since proteomics were done on bulk tissue), we found mild-moderate correlations between the datasets (**Figure S7D**). Overlapping genes with the *Camk2a* translatome were again mostly upregulated, whereas those with the *Pvalb* translatome were overwhelmingly downregulated (**Figure S7E**). When we compared specifically to proteomic studies of neocortex^50,52^, we found several synaptic genes among DEGs that overlapped with the *Camk2a* translatome (**Figure S7F-G**).

Finally, we compared our dataset with a transcriptomic study of human FXS fetal brain and patient-derived cerebral organoids^54^. We found an overlap between the FXS fetal brain transcriptome and 215 DEGs in the *Camk2a* translatome, as well as with 59 DEGs in the *Pvalb* translatome, but only the latter was significant (OR=1.47; **Figure S8A-B**). Amongst the 18 genes shared by all three, 8 were FMRP targets or ASD risk genes (**Figure S8C**). The overlap of DEGs between the FXS cerebral organoid transcriptome (at Day 84) and the *Camk2a* or *Pvalb* translatomes was not significant (**Figure S8D-F**). We found significant correlations between DEGs in human fetal brain and those in the *Camk2a* and *Pvalb* translatome (**Figure S8G**).

### *Rapgef4*: an ASD risk gene and FMRP target uniquely upregulated across cell types

While the initial goal of our study was to profile the translatome of excitatory and PV interneurons of *Fmr1* KO mice to provide insight on neuronal, circuit and behavioral phenotypes related to E/I imbalance, a second more translational goal was to find DEGs that could prioritize specific mechanisms for targeted therapy in FXS. Although several DEGs that were unique to either excitatory or inhibitory neurons were of interest, for example, because of their roles in synaptic assembly and function (**Figure 2**), the lack of pharmacological tools to target them and the paucity of research related to FXS did not make them attractive candidates. We also reasoned that genes similarly dysregulated across cell types would be more appropriate for translational purposes because drugs that targeted them would restore signaling pathways in both cell types.

We found 65 upregulated genes and 119 downregulated genes in both cell types, which were enriched for pathways associated with ‘Golgi membrane organization’, and ‘GTPase signal transduction’, respectively (**Figure 6A-B**). We found significant overlap with FMRP targets (27 DEGs; p=0.004), but not with ASD risk genes (12 DEGs; p=0.9), and a few (14 DEGs, p=0.9) were also enriched in the brain (see Methods; **Figure 6C**). Interestingly, only one gene, *Rapgef4* (*Epac2,* SFARI score=2)^55^, was at the intersection of all these categories. *Rapgef4* was significantly upregulated in both cell types and in both S1 and V1 of *Fmr1* KO mice (**Figure 6D**). EPAC2 (Exchange Protein directly Activated by cAMP 2) is a regulator of small GTPase signaling and a target of protein kinase A (PKA)-independent cAMP ^32,33,56–59^. Importantly, EPAC2 bidirectionally regulates synaptic stability and synaptic strength, such that upregulation of EPAC2 induces spine shrinkage, spine instability and less AMPAR content^32,60^, which are all known phenotypes in Fmr1 KO mice^61–65^.

**Figure 6.**
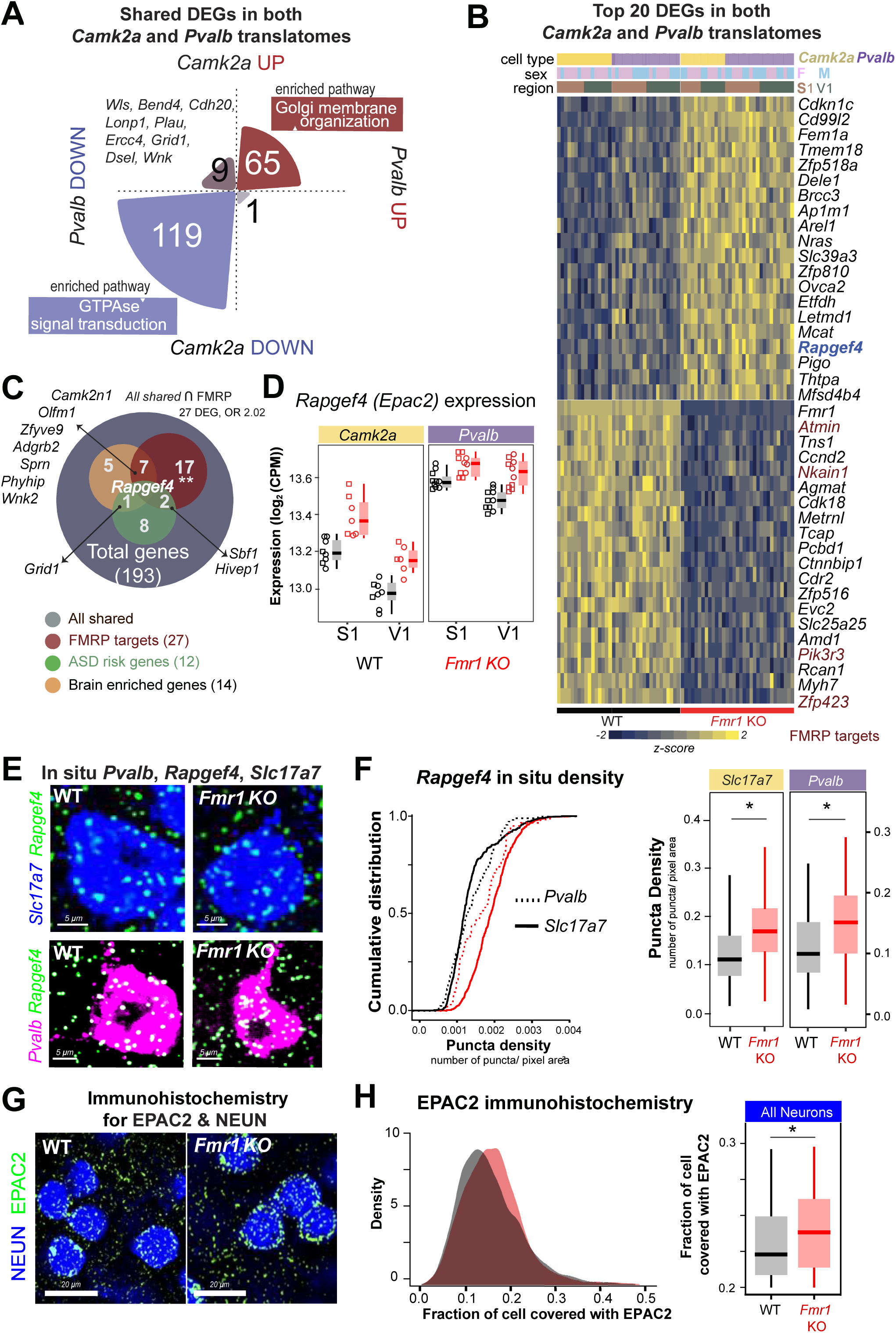
*Rapgef4* is a synaptic gene that is upregulated in both neuronal cell types in *Fmr1* KO mice and is also an ASD risk gene, an FMRP target, and enriched in the brain. **(a)** Shared DEGs between *Camk2a* and *Pvalb* translatomes, and corresponding GO enrichment terms for biological processes. **(b)** Heatmap of top 20 DEGs that share the same directionality in both cell types. FMRP target genes indicated in maroon. **(c)** Positive association between 193 shared DEGs (between *Camk2a* and *Pvalb* samples) and both FMRP targets (27 genes, OR=2.2, p=0.005; Fisher’s exact test) and ASD risk genes (12 genes; n.s.) Specific DEGs at major intersections are listed. **(d)** *Rapgef4* is significantly upregulated in both *Camk2a* and *Pvalb* samples across both S1 and V1 (FDR=0.05; *Pvalb*-S1: adj.p=0.048; *Pvalb*-V1: adj.p=0.01; *Camk2a*-S1: adj.p=0.002; *Camk2a*-V1: adj.p=0.008). **(e)** Example in situ hybridization probing for *Rapgef4*, *Pvalb* and *Slc17a7* (*Vglut1*) a marker of excitatory neurons. **(f)** Cumulative distribution (Left) and quantification of puncta density (Right) for *Rapgef4*+ puncta overlapping with *Slc17a7* or *Pvalb* in S1 of WT and *Fmr1* KO mice (*Slc17a7*: 0.17±0.002 vs. 0.12±0.001 puncta/pixel^2^, p<0.001; unpaired t-test; n= 2738/2009 neurons from n=3/4 WT and *Fmr1* KO mice, respectively; *Pvalb*: 0.14±0.004 vs. 0.10±0.003 puncta/pixel^2^, p<0.001; n=359/209 neurons from n=3/4 WT and *Fmr1* KO mice, respectively). Data represented as median±1.5*IQR. **(g)** Example immunohistochemistry for EPAC2 (green) and neuronal marker NEUN (blue). **(h)** Distribution (Left) of the fraction of cell soma area (NEUN+) covered by EPAC2 puncta in WT and *Fmr1* KO mice (WT: 0.126±0.004 vs. KO: 0.136±0.006; p=0.03; unpaired t-test; n=2618/2415 neurons from *n=*8/6 WT and *Fmr1* KO mice, respectively). *p < 0.05, **p < 0.01, ***p < 0.001

Transcripts of known protein interactors of EPAC2, including *Prickle2*, *Pdlim5*, *Kalrn*, *Rims1, Rims2, Unc13a* and *Unc13b*, showed higher expressions in *Fmr1* KO mice, suggesting a generalized upregulation of EPAC2 related pathways (**Figure S9A**). Seeded co-expression analysis identified several hundred genes that were positively or negatively correlated with *Rapgef4* expression, several of which were differentially expressed in *Fmr1* KO mice (**Figure S9B**). GO enrichment analysis of the correlated DEGs identified pathways known to be associated with EPAC2 and cAMP signaling, including ‘Protein exocytosis’^66^, ‘K^+^ ion transport’^67^, ‘Neuron death’^68^, ‘Autophagy’^69^ and ‘Ras signal transduction’^70^.

To validate the *Rapgef4* overexpression in *Fmr1* KO mice, we used multiplexed in-situ hybridization in S1 and demonstrated that excitatory and PV neurons had a significantly higher density of *Rapgef4* puncta in cortical tissue from *Fmr1* KO mice than in WT controls (**Figure 6E-F**; **Figure S9C**). Similarly, using immunocytochemistry, we found that EPAC2 puncta partially or completely overlapped with glutamic acid decarboxylase (GAD65), the vesicular glutamate transporter 1 (VGLUT1), and especially with the excitatory postsynaptic marker PSD-95 (**Figure S9D-E**). We further confirmed this synaptic localization of EPAC2 in Western blots of synaptoneurosomes from WT mice (**Figure S9F**). Using co-immunoprecipitation with antibodies against PSD95 or SYT2 (presynaptic inhibitory marker), we found EPAC2 was associated with both excitatory and inhibitory synapses (**Figure S9F-G**). EPAC2 puncta also overlapped with the neuronal marker NEUN, presumably representing perisomatic synapses (**Figure S9H**). The fraction of NEUN+ somata that was covered by EPAC2 staining was significantly higher in tissue from *Fmr1* KO mice (**Figure 6H**).

These results match those of previous studies showing that EPAC2 is localized to both postsynaptic dendritic spines, where it interacts with PSD-95 to regulate synaptic strength and plasticity^32,33^, and presynaptic axon terminals, where it regulates vesicle release and neurotransmission^60^. Considering how over-expression of EPAC2 in cortical neurons of WT mice leads to smaller, unstable spines that lack AMPA receptors^32^, i.e., known synaptic phenotypes of *Fmr1* KO mice^61–64,71,72^ it was reasonable to consider that EPAC2 would be a novel potential therapeutic target for FXS.

### Inhibition of EPAC2 using an antagonist ameliorates sensory and circuit phenotypes in *Fmr1* KO mice

To test whether the inhibition of EPAC2 can ameliorate certain sensory phenotypes of *Fmr1* KO mice we conducted calcium imaging and behavioral studies at P75-P110. Atypical sensory processing, including maladaptive defensive/avoidance behaviors and deficits in sensory/cognitive novel object recognition, have been described previously in FXS^19,43,73–77^. We used a well-characterized specific EPAC2 antagonist, ESI-05, which has previously been shown to have good bioavailability and CNS penetration^78,79^. Based on those studies, we administered ESI-05 to both WT and *Fmr1* KO mice using a dose of 10 mg/kg (i.p.) once daily for a period of 6 days.

We first tested whether ESI-05 could improve circuit phenotypes that are associated with sensory hypersensitivity^43^, namely a lower percentage of whisker responsive neurons in S1 and a decrease in the degree to which they adapt to repetitive whisker stimulation^19,27,43,76,80^. Using *in-vivo* 2-photon calcium imaging, we recorded from the same population of layer 2/3 neurons above the C2 barrel in *Fmr1* KO mice before and after chronic ESI-05 administration, or vehicle as a control (**Figure 7A-B**). ESI-05 treatment significantly increased the fraction of whisker-responsive neurons and their adaptation, whereas vehicle had no effect (**Figure 7C-E**).

**Figure 7:**
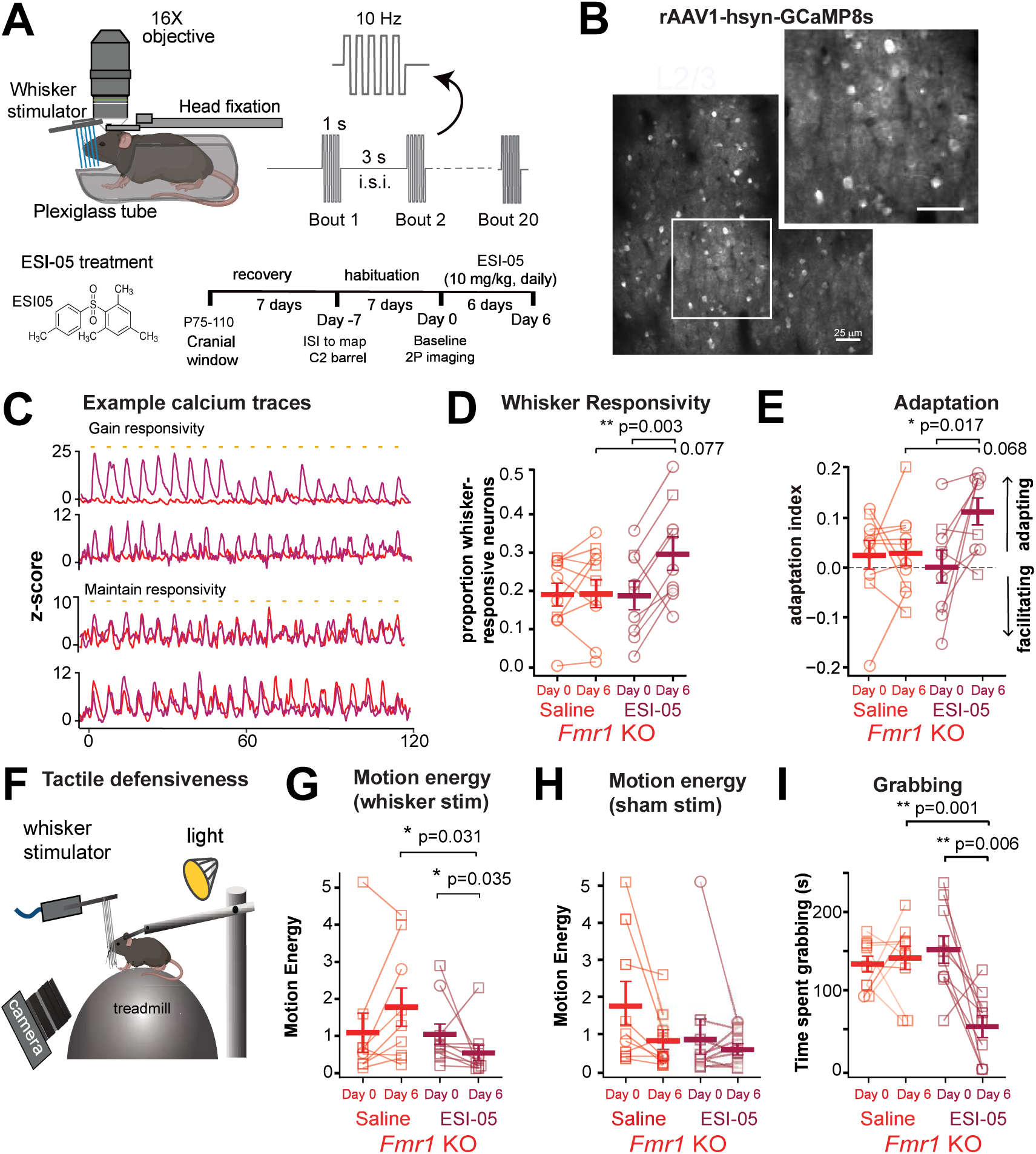
The EPAC2 antagonist ESI-05 ameliorates S1 circuit deficits and tactile defensiveness in adult *Fmr1* KO mice. **(a)** Top left: cartoon of set-up for 2P calcium imaging (2PCI) during whisker stimulation. Top right: trajectory of piezoactuator deflections for the 1 s stim duration, 3 s i.s.i. stimulation paradigm (10 Hz). Bottom left: Chemical structure of ESI05. Bottom right: experimental design and timeline of 2P imaging. **(b)** Representative FOV of GCaMP8s-expressing layer (L) 2/3 neurons above the C2 barrel. **(c)** Example traces of calcium transients in *Fmr1* KO mouse before (red trace) and after chronic ESI-05 treatment (purple trace). Orange tic marks indicate epochs of whisker stimulation. **(d)** Fraction of whisker-responsive Pyr neurons of *Fmr1* KO was significantly higher after 6 days of ESI-05 treatment compared to pre-treatment (ESI-05 Day 0 vs. Day 6: 0.12±0.04% vs. 0.30±0.03%, p=0.003), but saline treatment had no effect (Saline 0.19±0.03 vs. 0.19±0.03; p=0.9); day-treatment interaction p=0.018, n=10 and 9 mice, respectively. 2-way ANOVA with Bonferroni correction for d-e and g-i. **(e)** The adaptation index of Pyr neurons of *Fmr1* KO mice was higher after treatment with ESI-05 compared to pre-treatment (ESI-05 Day 0 vs. Day 6: 0.002±0.03 vs. 0.113±0.026; p=0.018), but saline had no effect (Saline: Day 0 vs Day 6, 0.024±0.025 vs. 0.026±0.029; p=0.9); time-treatment interaction p=0.042; time p=0.049). **(f)** Cartoon of tactile defensiveness behavior assay. **(g)** Motion energy (FaceMap) was significantly reduced in *Fmr1* KO mice after treatment with ESI-05 compared to pre-treatment (ESI-05 Day 0 vs. Day 6: 0.95±0.27 vs. 0.45±0.21 a.u., p=0.03), but not in saline controls (Saline Day 0 vs. Day 6: 0.99±0.53 vs. 1.68±0.51 a.u); time-treatment interaction p=0.014. **(h)** Same as *H* but for sham stimulation where the whisker stimulator does not contact the whiskers. There was no significant effect of ESI-05 treatment, although mice treated with saline showed behavioral habituation with less motion at Day 6 (Saline: 0.74±2.60 vs. 1.80±0.24 a.u., p=0.034; ESI-05: 0.49±0.14 vs. 0.83±0.50 a.u., p=0.95). **(i)** Duration of time spent grabbing the stimulator was significantly reduced in *Fmr1* KO mice after treatment with ESI-05 compared to pre-treatment (ESI-05 Day 0 vs. Day 6: 148±0.17 vs. 52±0.14 s; p=0.0001), but not in vehicle controls (Saline Day 0 vs. Day 6: 130±0.1 vs. 138±0.14 s), time-treatment interaction p=0.0007). *p < 0.05, **p < 0.01, ***p < 0.001

Next, we used a behavioral assay (**Figure 7F**) in which *Fmr1* KO show significantly more avoidant and defensive behaviors than WT mice to repetitive whisker stimulation^19,43,76^. Using *Facemap*^81^ to analyze videos of mice, we found that *Fmr1* KO mice manifested a significant reduction in motion energy after treatment with ESI-05 (n=10), whereas those treated with vehicle (n=9) showed no difference (**Figure 7G**). These effects were specific to stimulation because under sham stimulation we observed no differences in motion energy (**Figure 7H**). We also observed a significant decrease in the amount of time that *Fmr1* KO mice spent grabbing the stimulator after ESI-05 treatment, but not after vehicle treatment (**Figure 7I**).

*Fmr1* KO mice also show deficits in the novel object recognition task (NORT)^73–75,77^. We used NORT to test two sensory modalities, tactile (rough vs. smooth textures) and visual (yellow vs. black colors) (**Figure 8A**). We tracked the movements of WT or *Fmr1* KO mice with *DeepLabCut*^82,83^ and calculated how much time they spent around novel or familiar objects. Treatment with ESI-05 for 6 days had no impact on both the total distance explored by mice or their patterns of exploration for either genotype (**Figure 8C, G**). The same was true in the open field test, where mice of both genotypes showed a similar preference for exploring the edges and corners and avoided the center of the arena (**Figure S10**). We then habituated mice to two objects of similar size, shape and texture for a period of 10 min and found no differences in how much time WT and *Fmr1* KO mice spent interacting with these objects (not shown). Next, one of the objects was randomly replaced with a novel object of a different texture or color. WT mice (regardless of treatment) had a clear preference for the novel texture; however, whereas vehicle-treated *Fmr1* KO showed no such preference, ESI-05 treatment led to a significant improvement in their novelty-seeking behavior (**Figure 8B, D**). Similarly, in the color discrimination version of NORT, whereas *Fmr1* KO mice tended to avoid the novel color object, this was also rescued by ESI-05 treatment and their behavior was indistinguishable from WT controls (**Figure 8F, H**). Crucially, this effect of ESI-05 was specific to *Fmr1* KO mice, as no differences were detected in WT controls treated with ESI-05 or vehicle. Thus, an established inhibitor of EPAC2 was highly efficacious in preclinical studies aimed at restoring circuit and behavioral phenotypes related to sensory processing in *Fmr1* KO mice.

**Figure 8:**
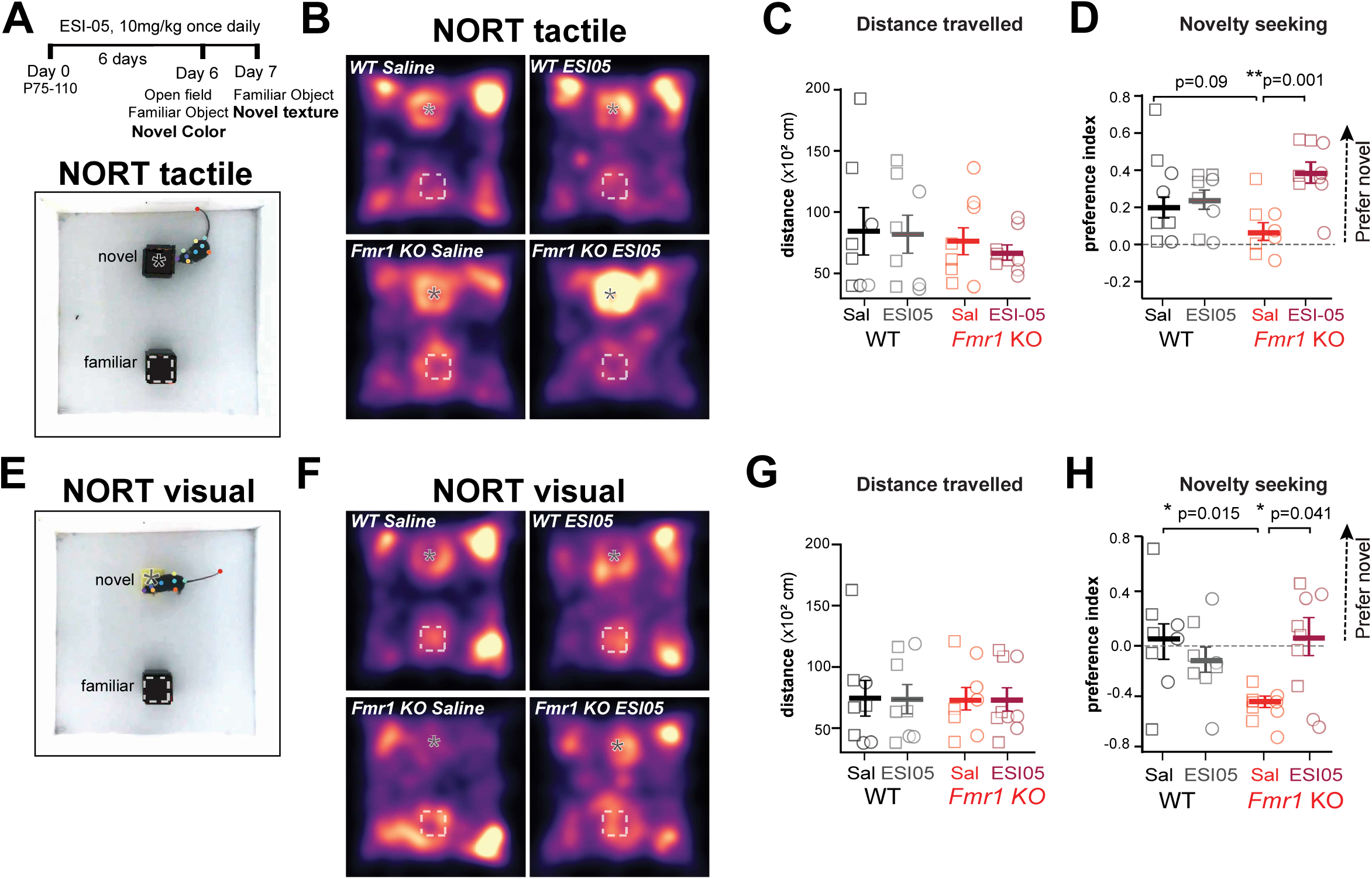
ESI-05 improves novel object recognition by *Fmr1* KO mice. **(a)** Top: Experimental design and timeline of behavioral assessments. Bottom: representative frame of movie recording NORT-tactile; note DLC labels tracking mouse. **(b)** Mean heatmaps of exploration during NORT-tactile for WT and *Fmr1* KO mice treated with saline (8 WT and 8 KO mice) or ESI-05 (8 WT and 9 KO mice). **(c)** Distance travelled was similar across all groups (p>0.05, 2-way ANOVA with Bonferroni correction for panels C-D and G-H). **(d)** The preference index for a novel object of a different texture (based on the time spent investigating each object) was significantly higher after ESI-05 treatment in *Fmr1* KO mice (saline vs. ESI-05, p=0.001), but not in WT controls (p=0.9). **(e)** Representative frame of movie recording NORT-visual. **(f)** Mean heatmaps of exploration during NORT-visual for WT and *Fmr1* KO mice treated with saline or ESI-05 (same # mice as in *B*). **(g)** Distance travelled was similar across all groups. **(h)** The preference index for a novel object of a different color was significantly lower in saline-treated *Fmr1* KO mice than in WT mice (p=0.03). ESI-05 treatment significantly increased the preference index in *Fmr1* KO mice (p=0.04), but not in WT mice (p=0.85). *p < 0.05, **p < 0.01, ***p < 0.001

## DISCUSSION

Here, we profiled the translatome of excitatory and PV inhibitory neurons in *Fmr1* KO mice, the principal model of FXS. We found that translatome dysregulation is strikingly different in these cell types, with excitatory neurons showing >5-fold more DEGs than PV neurons. Moreover, the signaling pathways affected were different (often opposite) between cell types. Amongst 184 DEGs that were concordantly dysregulated in excitatory and inhibitory populations, we identified *Rapgef4*, as the only one that was a direct target of FMRP, an ASD risk gene, and enriched in the brain. When overexpressed in cortical neurons, the protein it encodes, EPAC2, is known to reduce the size, turnover, and AMPA-receptor content of dendritic spines, which happen to be synaptic phenotypes observed repeatedly in *Fmr1* KO mice. In preclinical studies we discovered that chronic treatment of *Fmr1* KO mice with a specific inhibitor of EPAC2 can fully rescue circuit and behavioral phenotypes related to atypical sensory processing in S1. We conclude that EPAC2 is a potential novel target for therapy in FXS.

This was the first attempt to directly compare the translatome of excitatory and inhibitory neurons in a mouse model of ASD. We were motivated by the debate that currently exists around the true culprit of E/I imbalance in ASD/FXS^80^: which are more impacted, excitatory or inhibitory neurons? A priori it might have been expected that loss of FMRP, with its essential roles in cellular translation, would affect all neurons in similar ways. Alternatively, loss of FMRP could have unique effects on different cell types, if only to explain the observed E/I imbalance. We found a remarkable degree of cell-specific translatome dysregulation, with excitatory neurons bearing the brunt of the loss of FMRP. One possible reason is that *Camk2a* neurons represent a much more heterogeneous population (including intratelencephalic, pyramidal tract projecting, or layer 4 spiny stellate neurons) than PV-expressing interneurons. Interestingly, although roughly equal numbers of genes in *Camk2a* neurons were up- and downregulated in *Fmr1* KO mice, DEGs in *Pvalb* neurons were overwhelmingly downregulated.

The signaling pathways affected are also different. In excitatory neurons, chromatin remodeling’, ‘autophagy’/’protein catabolism’, and ‘synapse organization’ were all upregulated, which makes sense based on what we know about FMRP’s role in translational suppression of key synaptic genes^5,36,84^. Indeed, we identified *Setd2* among several other chromatin modifying genes that were upregulated in excitatory cells **(Figure S5A)**, as previously reported^36^. WGCNA also identified ‘histone modification’ and ‘chromatin remodeling’ as top enriched pathways in the *Camk2a*-specific black module. In contrast, most DEGs in *Pvalb* neurons were downregulated and tended to be related to neuronal development: ‘cell growth’, ‘synapse organization’, and ‘axonogenesis’. This was also reflected in the WGCNA’s lightgreen module, which was specifically downregulated in PV neurons and was enriched in pathways like ‘axonogenesis’, neuron differentiation’ and ‘proteoglycan synthesis. Even for the significant overlap between FMRP targets or ASD risk genes and unique DEGs in either *Camk2a* or PV neurons, the affected pathways were once again distinct for each cell type, and in some cases, dysregulated in opposite ways. Overall, excitatory neurons in *Fmr1* KO mice show upregulation of chromatin remodeling, protein catabolism, and synapse organization pathways, while PV interneurons demonstrated downregulation of neuronal/synaptic maturation and function pathways, which fits with the notion that PV neurons are hypofunctional in FXS, as previously reported^19,21,22,26^.

Symptoms of FXS, ASD, epilepsy, and other NDCs, have been proposed to arise from an imbalance in the ratio of excitation to inhibition (E/I) in the brain^20^. While this theory provides a compelling framework for understanding how certain symptoms (e.g., seizures, sensory hypersensitivity) could arise from excess neuronal activity, ultimately such complex diseases cannot be explained by a unidimensional model but reflect instead more nuanced changes in both cell types^26,27,85^. Based on our RNA-seq results, we surmise that unique changes in excitatory and inhibitory neuron physiology in FXS, which contribute to E/I imbalance, arise from the profound cell type specific transcriptional differences. But this is also surprising because mutations in *Fmr1* or in other ASD-associated genes that are expressed in both excitatory and inhibitory neurons (e.g., synaptic genes like *Syngap1*, *Nrxn1*, *Cntnap2*) could have affected them similarly. It underscores the importance of our study and raises new questions. Developmentally, when do these differences in gene expression first arise? It has been proposed that hypofunction of PV neurons, which is apparent very early postnatally in *Fmr1* KO mice^43^, could represent a final common pathway for many NDCs^22^, but this could also reflect compensation for a primary defect in excitatory neurons^27^. Moreover, the fact that fewer genes are differentially expressed in adult PV neurons than in MGE interneurons at P15 means there could be some adaptive compensation by inhibitory neurons. We also found significant overlap and correlation between DEGs in our dataset and those identified in fetal human brain and cerebral organoids derived from neurons of FXS patients, and many of the shared DEGs were targets of FMRP, ASD risk genes, or both. Only a developmental study that examines transcriptional and/or proteomic dysregulation across cell types from embryonic to adult stages could directly resolve this. It also remains to be determined whether similar unique cell-specific gene dysregulation is triggered by other ASD-associated mutations.

Despite the dramatic cell type-specific differences in translatome dysregulation, there were also nearly 200 DEGs that were shared between excitatory and inhibitory neurons, as well as some shared biological pathways, such as the downregulation of ‘axonogenesis’ and ‘Ras/GTPase signal transduction’. Thus, FMRP does play some conserved roles across cell types. Importantly, this analysis led to the identification of *Rapgef4*/EPAC2 upregulation as an attractive target for therapy based on its known roles in synaptic development and plasticity^32,33^. Expression levels of its binding partners *(Prickle2, Dlgap1, Kalrn*, *Rims1*, *Rims2*, *Pclo*, *Bsn Unc13a*, *Unc13b*) are also higher in *Fmr1* KO mice **(Figure A9A-B)**, and several are known to be upregulated at the protein level in *Fmr1* KO mice at P17 (e.g., Prickle2, Piccolo, Bassoon)^50,52^. Furthermore, pathways involving MAPK and RAS signaling were also upregulated (not shown), and this fits with published data showing that MAPK1, which is downstream of RAP1 (a target of EPAC2), is also overexpressed in the hippocampal proteome at P17^50,52^.

*Rapgef4* had not been identified as a DEG in our study of MGE cortical interneurons at P15, and EPAC2 was not significantly dysregulated in proteomic studies of the neocortex at P17^50,52^ or the hippocampus at P15^51^. Moreover, *Rapgef4* was significantly downregulated in organoids from iPSCs of FXS cases **(Figure 8)**. This suggests that *Rapgef4*/EPAC2 upregulation in FXS may emerge gradually during brain development (P15 in mice roughly corresponds to birth in humans), making it an ideal candidate for adult intervention. Future studies will need to address whether EPAC2 is also elevated at synapses in post-mortem human tissue from adult cases of FXS.

Perhaps the most exciting result was that treatment with an EPAC2 antagonist can rescue circuit phenotypes and several behavioral phenotypes in Fragile X mice without any observed adverse effects. Unlike EPAC1, which is expressed ubiquitously in the body, EPAC2 is predominantly expressed in the brain and to a lesser extent in other neuroendocrine tissues. This bodes well for future preclinical trials and safety studies in humans, as compounds that target EPAC2 should not have effects outside the CNS. Our results should encourage the development of novel EPAC2 inhibitors for the treatment of FXS. More generally, our study exemplifies how transcriptomic approaches in animal models of neuropsychiatric conditions can be used to prioritize potential novel therapeutic targets.

## Supplemental Figures

**Figure S1 (Related to Figure 1):**
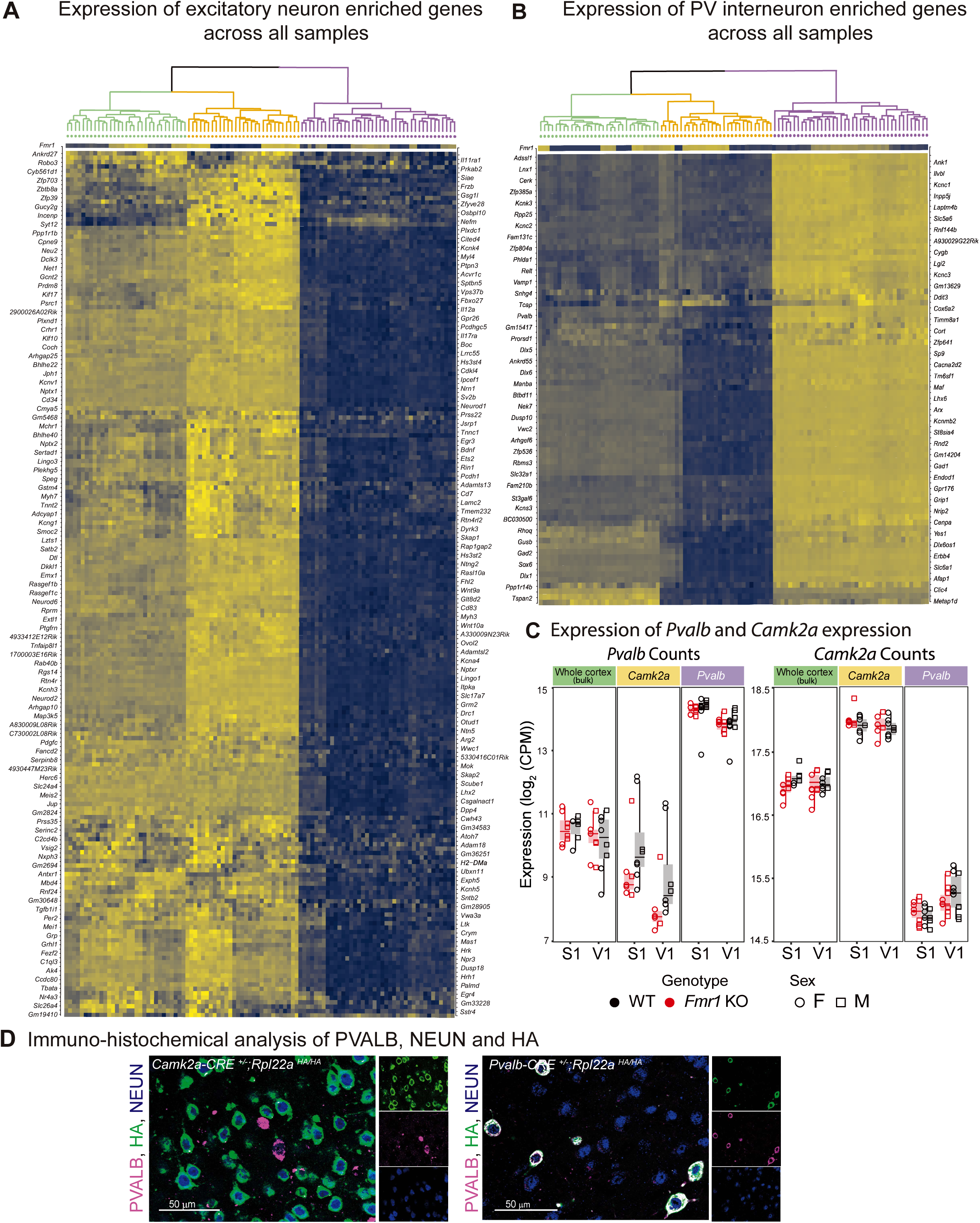
Cell type specificity of RiboTag approach. **(a, B)** The *Camk2a*-specific (A) and *Pvalb*-specific (B) translatomes are enriched for excitatory and inhibitory neuron transcripts, respectively. The list of cell type-specific genes was generated by an unbiased comparison with the Allen Brain single-cell dataset (see Methods). **(C)** *Pvalb* and *Camk2a* gene expression levels were higher in their respective samples, showing specificity of the RiboTag approach. **(D)** Representative immunohistochemistry images through S1 using antibodies against PVALB, NEUN, and the HA Tag. Note overlap in expression of PVALB and HA in *Pvalb*-Cre^+/−^;*Rpl22a^HA/HA^* mice, but not *Camk2a*-Cre^+/−^;Rpl22a^HA/HA^ mice.

**Figure S2 (Related to Figure 2C):**
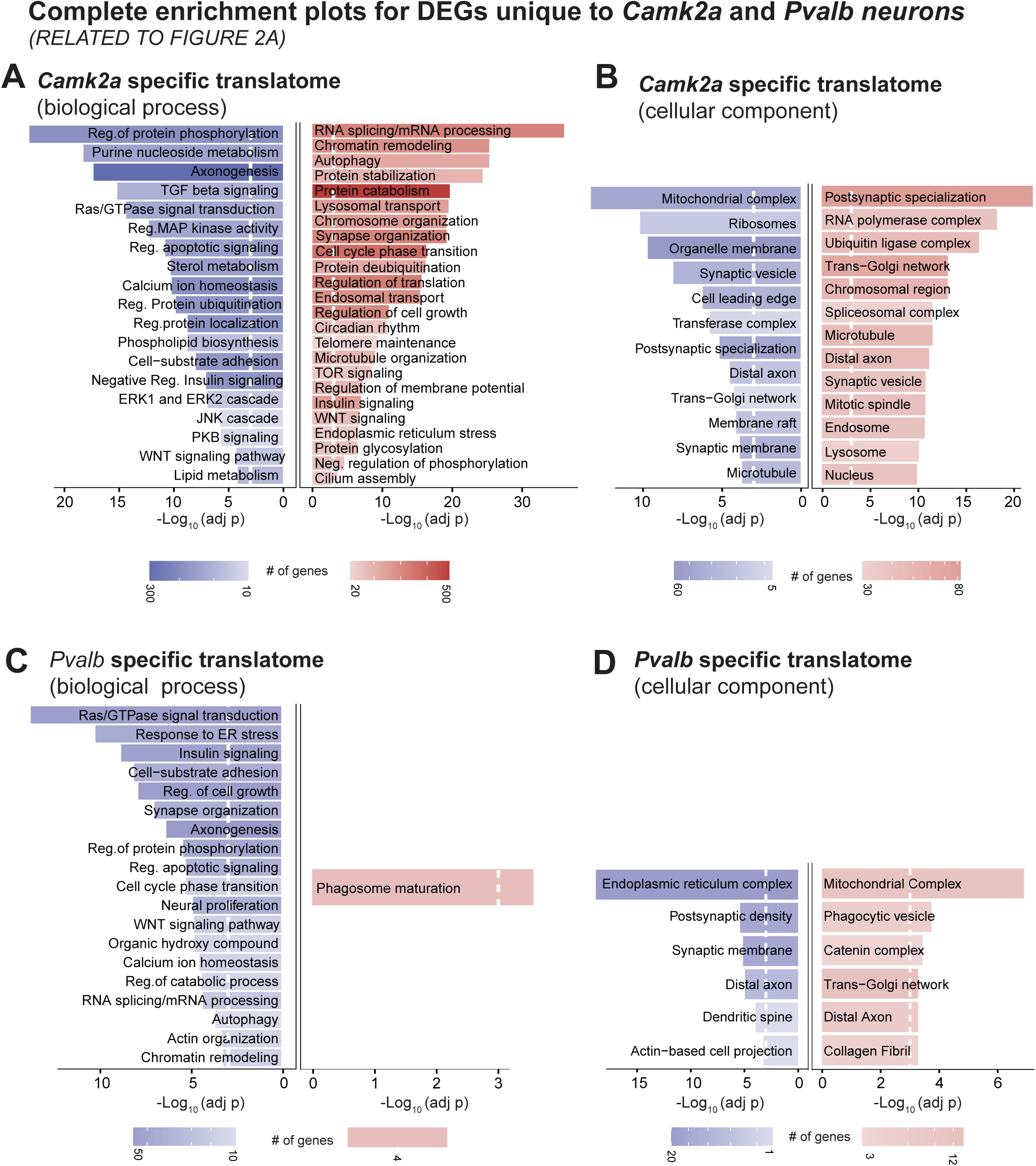
GO enrichment pathways in *Camk2a* and *Pvalb* translatomes. **(a)** GO enrichment terms for biological process in *Camk2a*-unique DEGs. White line indicates threshold of significance (FDR=0.05; p.adj<0.05). Terms collapsed based on similarity (see Methods). GO enrichment terms of upregulated genes indicated in red and downregulated genes in blue for all Figures **(B)** Same as *A* but GO terms for cellular component. **(C)** Same as *A* but for *Pvalb*-unique DEGs. **(D)** Same as *B* but for *Pvalb*-unique DEGs.

**Figure S3 (Related to Figure 2C):**
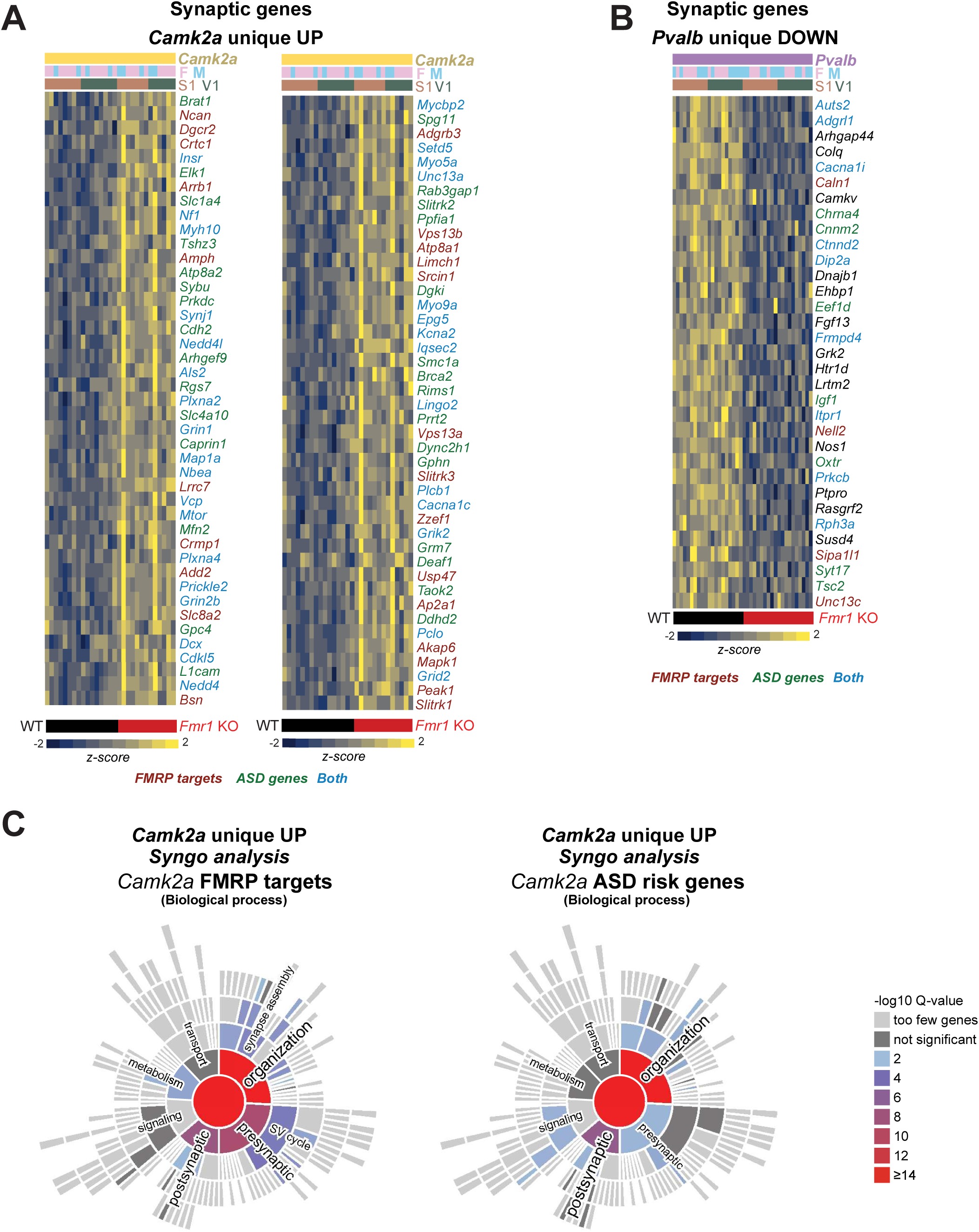
Synaptic DEG’s of unique *Camk2a Up a*nd *Pvalb* DOWN DEGs. **(a)** Heatmap of *Camk2a* unique synaptic genes that were also FMRP target and ASD risk genes. **(B)** Heatmap of *Pvalb* unique synaptic genes including FMRP target and ASD risk genes. **(C)** Syngo enrichment analysis of the FMRP and ASD risk genes among the *Camk2a* DEGs.

**Figure S4 (Related to Figure 2C):**
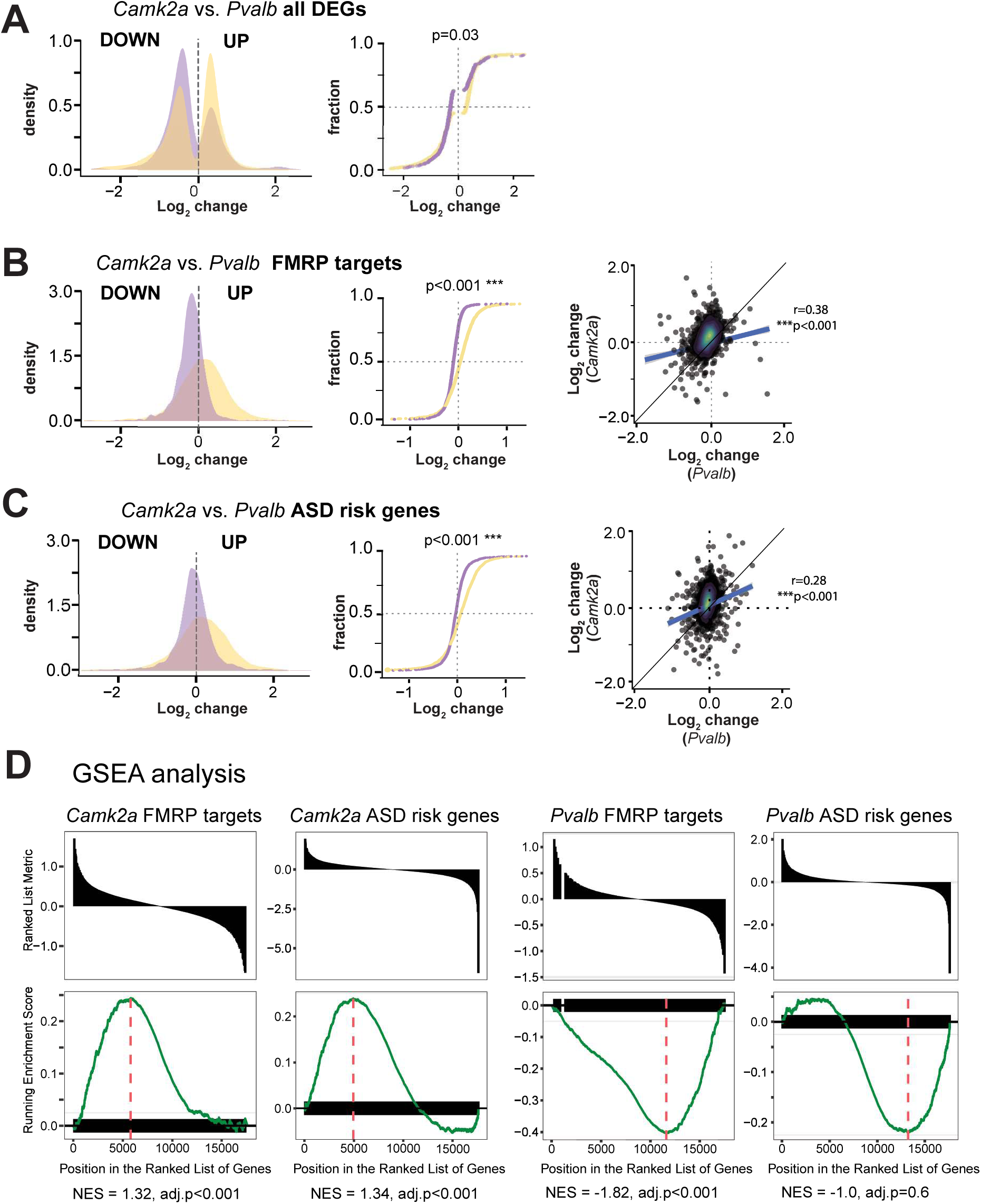
FMRP target and ASD risk genes among *Camk2a* unique DEGs are mostly upregulated whereas those among *Pvalb* unique DEGs are mostly downregulated. **(a)** (left) Density distribution of log_2_ fold change for all *Camk2a* and *Pvalb* DEGs. (right) Cumulative distribution of *Camk2a* and *Pvalb* DEGs shows a significant difference, with *Camk2a* unique DEGs generally being upregulated in *Fmr1* KO mice, while *Pvalb* unique DEGs are downregulated (p=0.03; Wilcoxon Rank Sum and Signed Rank Test). **(B)** (left) Density distribution of log_2_ fold change for all FMRP targets (including non DEGs) for *Camk2a* and *Pvalb* translatomes. (center) Cumulative distribution of *Camk2a* and *Pvalb* FMRP targets shows significant difference (p<0.001). (right) Scatter plot comparison of log2 fold change of all FMRP targets in *Camk2a* and *Pvalb* translatomes (rho=0.39, p<0.001, Spearman). **(C)** Same as *B* but for ASD risk genes. p<0.001 for center panel, and rho=0.28, p<0.001 for right panel. **(D)** GSEA analysis of *Camk2a* and *Pvalb* DEGs that are also FMRP target and ASD risk genes. *Camk2a* FMRP: adj.p=9.37e-8, NES=1.89; *Camk2a* ASD: adj.p=7.23e-7, NES=1.70; *Pvalb* FMRP: adj.p=0.146, NES=-1.28; *Pvalb* ASD: adj.p=0.54, NES=-0.95. *p<0.05, **p<0.01, ***p<0.001

**Figure S5 (Related to Figure 3):**
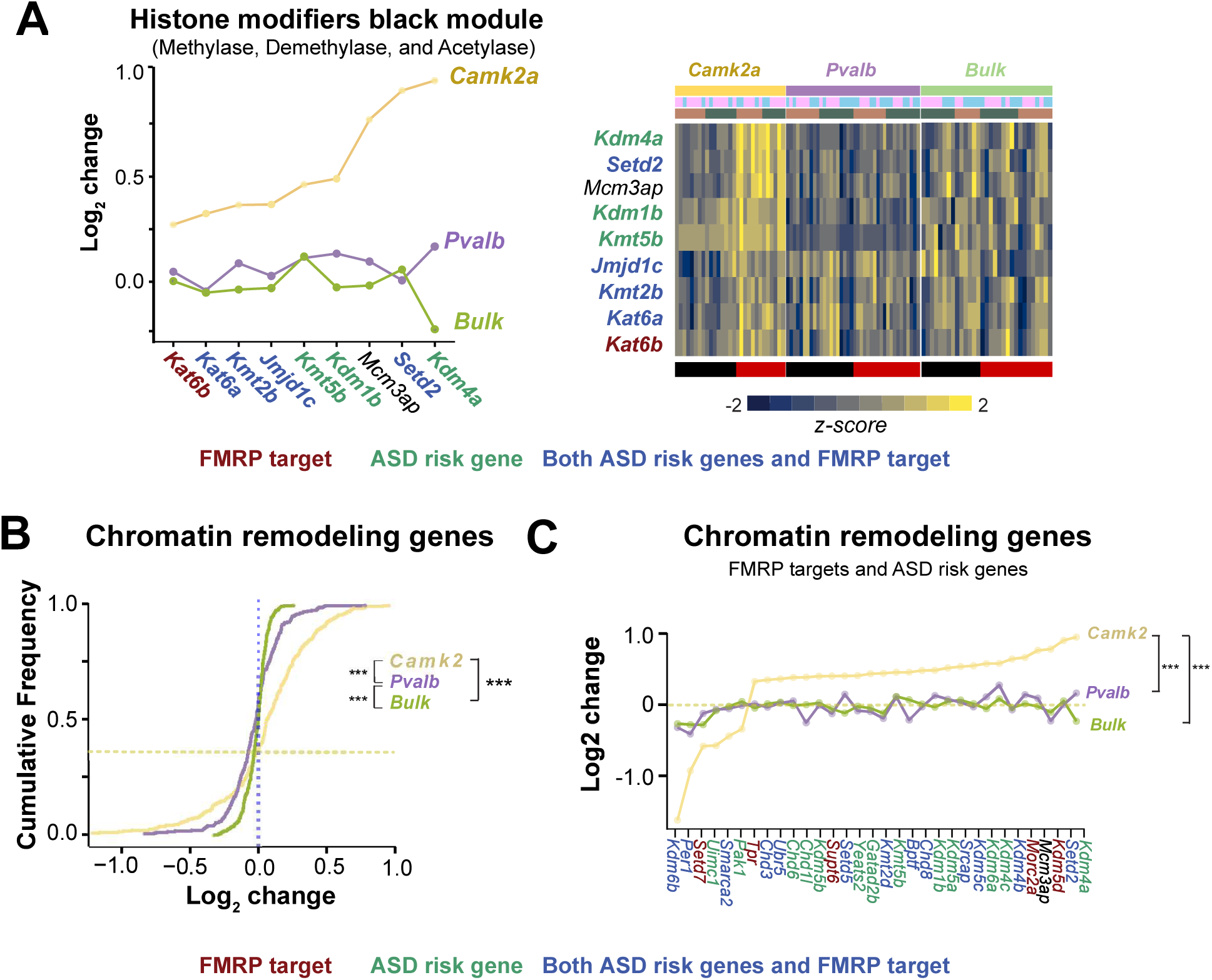
Chromatin genes that are also FMRP targets and ASD risk genes are upregulated in the *Camk2a* translatome of *Fmr1* KO mice. **(a)** (left) Log_2_ fold change of histone modifiers (only methylase, demethylase and acetylase genes shown) from the black module for *Camk2a*, *Pvalb* and bulk samples. Most of the genes are FMRP targets, ASD risk genes, or both. (right) Heatmap of the corresponding genes. **(b)** Cumulative frequency distribution of log_2_ fold-change values for all chromatin remodeling genes across *Camk2*, *Pvalb*, and bulk samples. DEGs in *Camk2a* samples were significantly upregulated in *Fmr1* KO mice (*Camk2a* vs *Pvalb*: p=5.5e-10; *Camk2* vs. bulk: p=6.6e-16; Kolmogorov-Smirnov test with Bonferroni correction for multiple comparisons). The difference between *Pvalb* and bulk samples was also significant (p=4.71e-4). The yellow horizontal line indicates the frequency at which the *Camk2* genes are upregulated (0.4). **(c)** Chromatin remodeling genes that are also ASD risk genes, FMRP targets, or both, were upregulated in the *Camk2* of *Fmr1* KO mice (yellow), relative to the *Pvalb* (purple) or bulk (green) transcriptomes (*Camk2a* vs *Pvalb*: p= 2.75e-10; *Camk2a* vs. bulk: p=*6.600e-16*, *Pvalb vs bulk: p= 0.9,* Kolmogorov-Smirnov test with Bonferroni correction for multiple comparisons )

**Figure S6 (relates to Figure 4):**
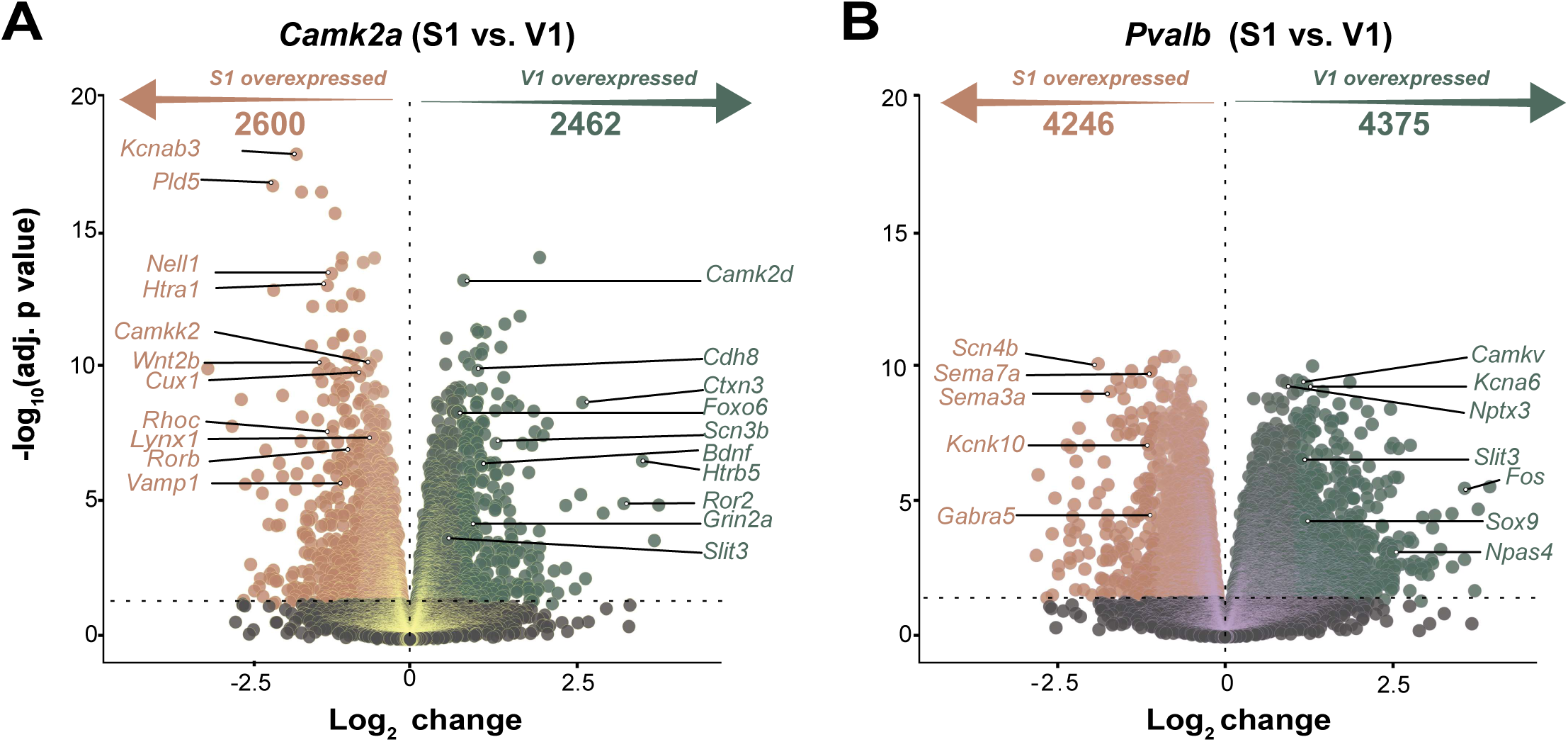
The translatomes of S1 and V1 are distinct from each other in WT mice. **(a)** Volcano plots showing which genes in the *Camk2a* excitatory translatome of WT mice are differentially expressed in S1 vs. V1. 2,462 genes were overexpressed in V1 and 2,600 were overexpressed in S1, respectively. Horizontal dotted line marks the threshold of significance adjusted p<0.05 (log_10_). **(B)** Same as *A* but for *Pvalb* inhibitory neuron translatome. 4,375 genes are overexpressed in V1, and 4,246 genes are overexpressed in S1.

**Figure S7 (Related to Figure 5):**
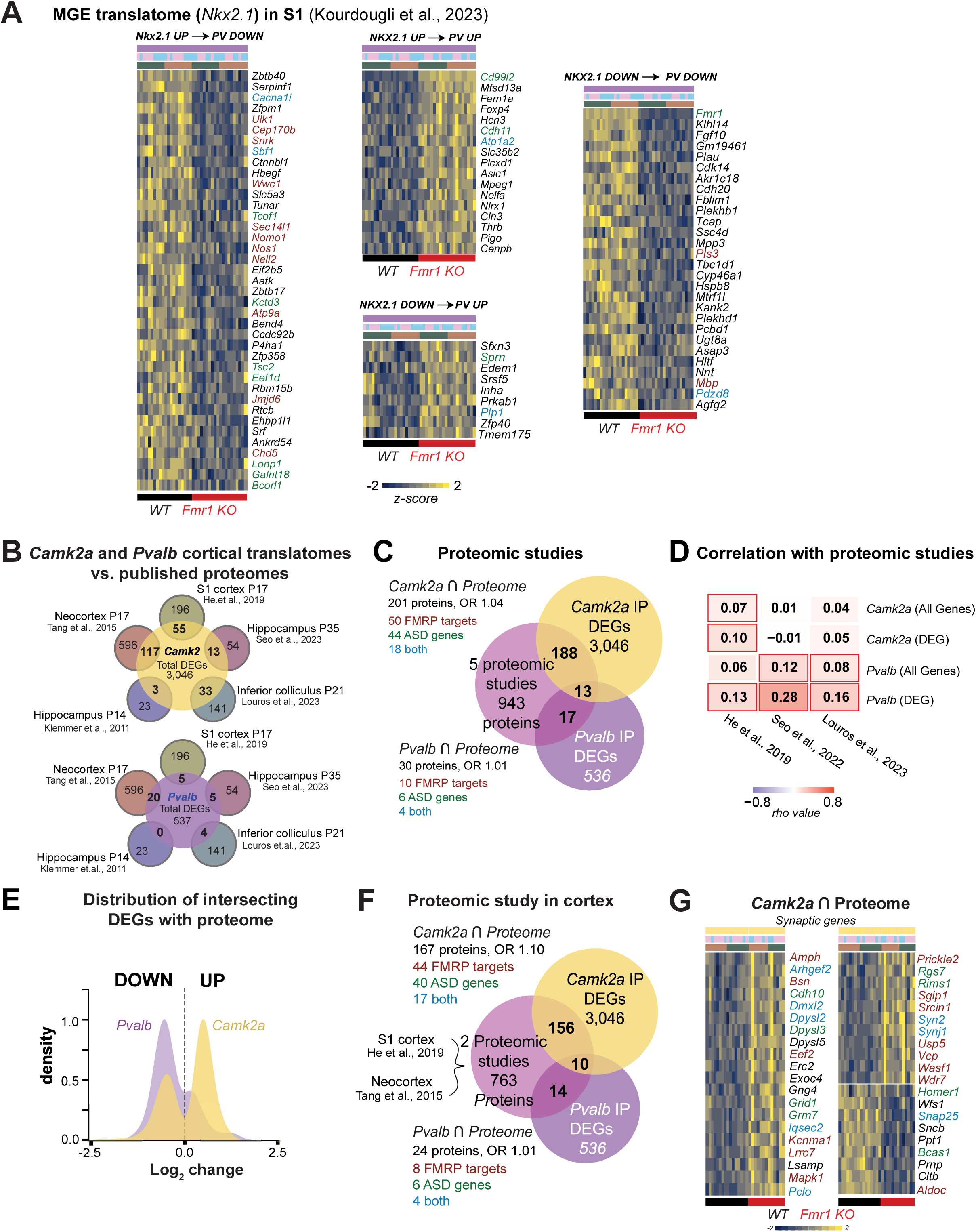
Comparison with published MGE translatome and proteomal data of *Fmr1* KO mice. (A) Heatmap of DEGs in *Nkx2.1* translatome at P15 that maintain or lose directionality of dysregulation in *Pvalb* transcriptome. Genes that are FMRP targets, ASD risk genes, or both are indicated. (B) Overlaps between *Camk2a* and *Pvalb* cortical translatomes and five published proteomes from *Fmr1* KO mice. (C) Comparison of *Camk2a* and *Pvalb* DEGs with a combined set of 943 differentially expressed proteins from published cortical proteomes (showed no significant associations (*Camk2a*∩proteomes: p=0.65; *Pvalb*∩proteomes: p=0.9). (D) Heatmap of synaptic genes that are both *Camk2a* DEG and combined proteome DEG. (E) Frequency distribution of fold change of DEGs shared between proteomic data and *Camk2a* (201 genes) and *Pvalb* translatomes (30 genes). (F) Correlation coefficients between the DEG proteomes and either *Camk2a* or *Pvalb* translatomes (all genes or only DEGs). Red outlines indicate significant correlation.

**Figure S8 (Related to Figure 5):**
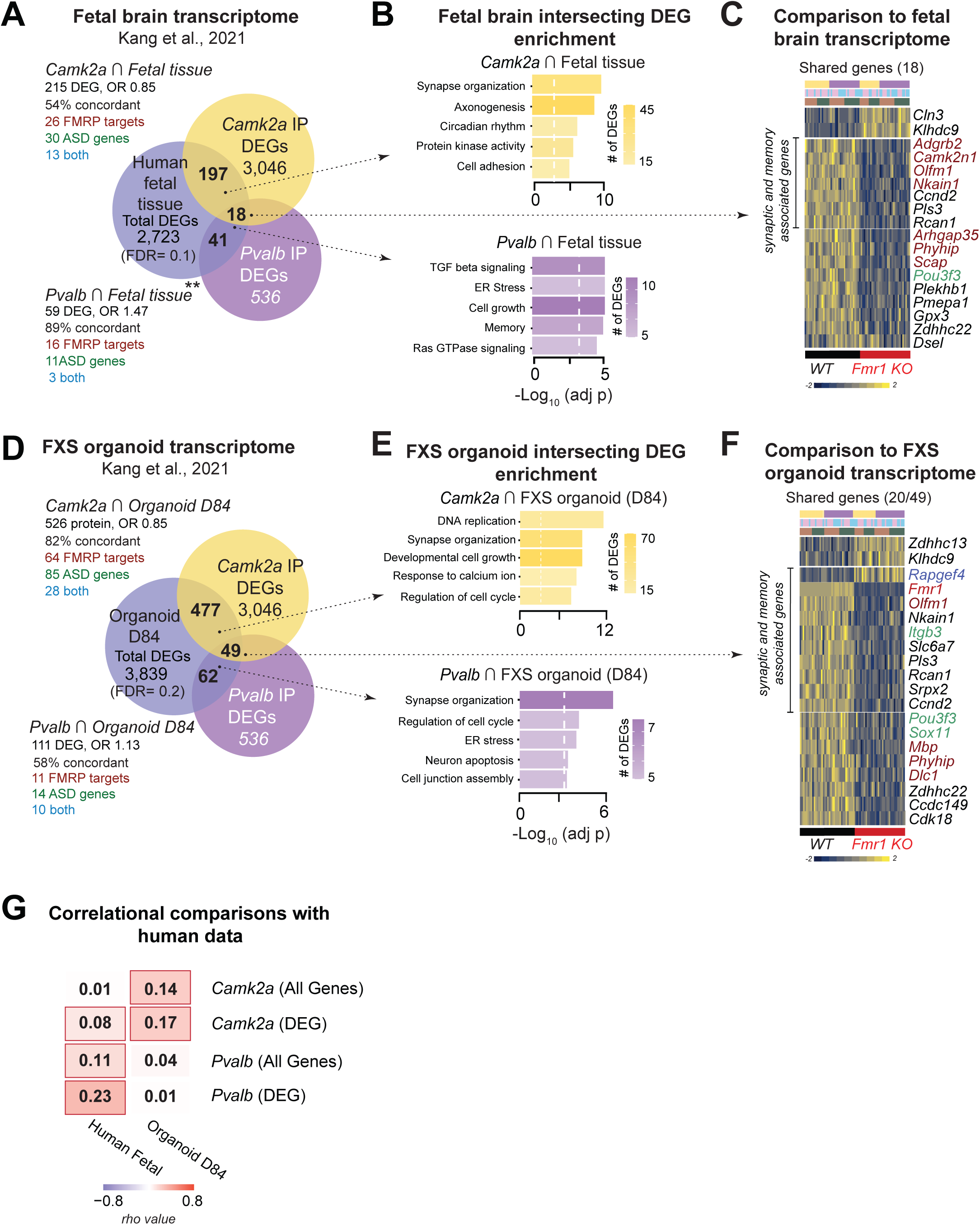
Comparisons with published proteomes of Fmr1 KO mice and transcriptome of FXS-derived human neurons. (A) Significant and positive association between DEGs (*Fmr1* KO vs WT) from adult *Pvalb* translatome (but not the *Camk2a* translatome) and DEGs from fetal human transcriptome (Kang et al., 2021) (OR=1.47, p=0.008). (B) Go enrichment terms of DEG’s from *Camk2a a*nd *Pvalb* translatomes with DEG’s from fetal FXS transcriptome. White line marks −log_10_ p-adj threshold of 0.05. (C) Heatmap of all DEGs between *Camk2a* and *Pvalb* translatomes that overlap with human FXS fetal brain DEG (from Kang et al. 2021). Synaptic and memory associated genes, as identified by enrichment analysis, are indicated. (D) Comparisons of DEG’s from *Camk2a* and *Pvalb* translatomes with DEG’s from FXS patient IPSC-derived organoids (D84) (from Kang et al. 2021). (E) Go enrichment terms of *Camk2a* and *Pvalb* DEG ∩ human FXS organoids DEG (D84). White line marks −log_10_ p-adj threshold of 0.05. (F) Heatmap of all DEGs between *Camk2a* and *Pvalb* translatomes that overlap with FXS patient IPSC-derived organoids DEG (from Kang et al. 2021). Synaptic and memory associated genes, as identified by enrichment analysis, are indicated. (G) Correlation coefficients between Fetal and Organoid (D84) transcriptome with either *Camk2a* or *Pvalb* translatomes (all genes or only DEGs).

**Figure S9 (Related to Figure 6):**
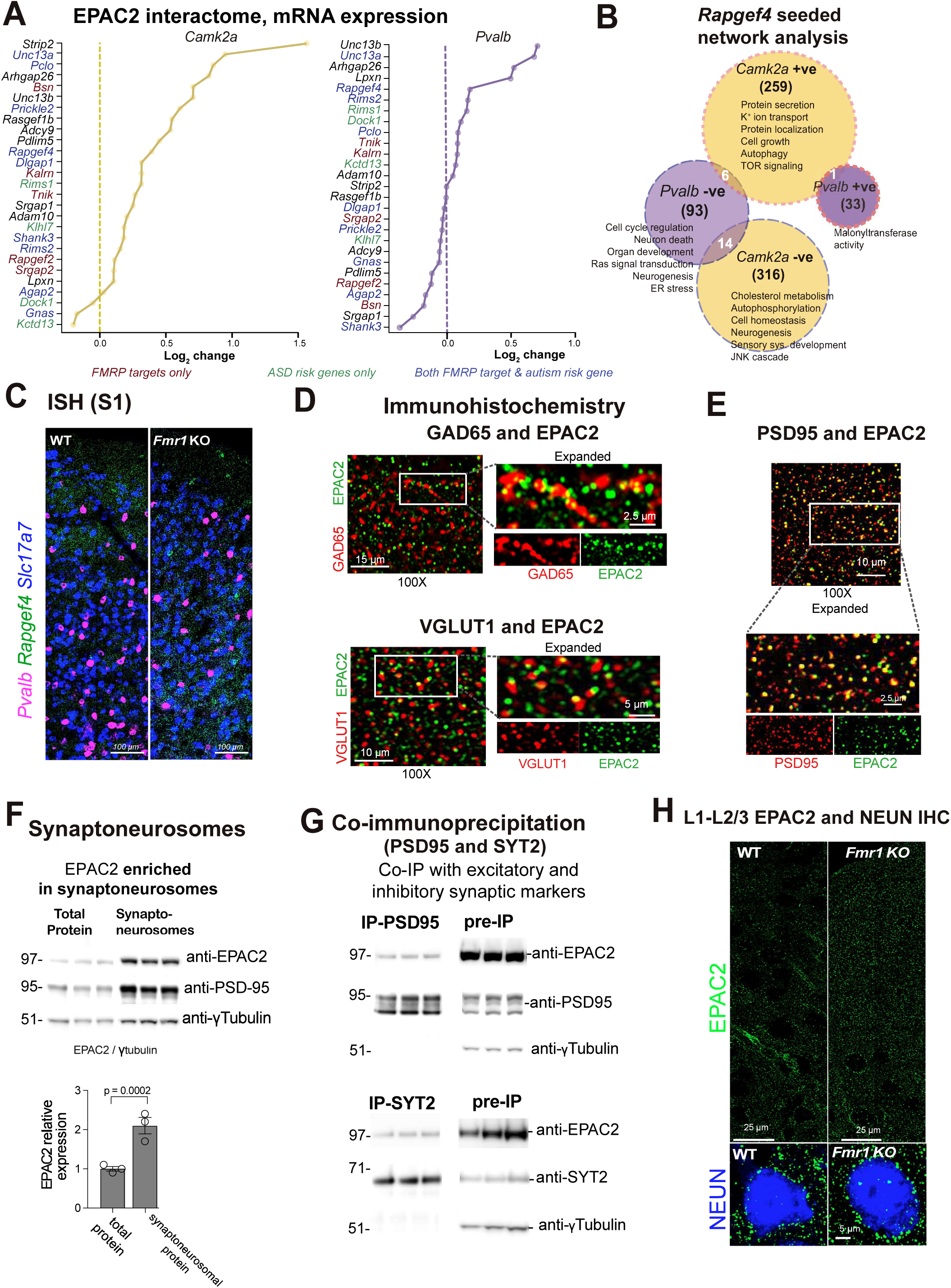
EPAC2 is enriched at synapses. **(a)** While most genes in the EPAC2 interactome were upregulated in *Camk2a* samples from *Fmr1* KO mice, their dysregulation was more balanced in *Pvalb* samples. Note that most of these genes are also FMRP targets, ASD risk genes, or both. **(B)** Seeded network analysis of *Rapgef4* with corresponding pathways of correlated and anticorrelated genes. **(C)** Representative in situ hybridization image for expression of *Rapgef4*, *Pvalb*, *Slc17a7* in S1. **(D)** Colocalization of GAD65 (top) and VGLUT1 (bottom) (in red) with EPAC2 (green). **(E)** Colocalization of PSD95 (red) with EPAC2 (green). **(F)** (Top) Western blot showing enrichment of EPAC2 in synaptoneurosomal preparations. (Bottom) EPAC2 levels are higher in the synaptoneurosomal fraction compared to the total lysate (p=0.0002, t-test). **(G)** Western blots showing co-immunoprecipitation of EPAC2 with PSD95 (top) and SYT2 (bottom). Note that the intensity of the bands cannot be compared across the two experiments as the quantities loaded differed between IP-PSD95 and pre-IP. **(H)** Representative image of Epac2 and NEUN (Related to Figure 6G).

**Figure S10 (Related to Figure 8):**
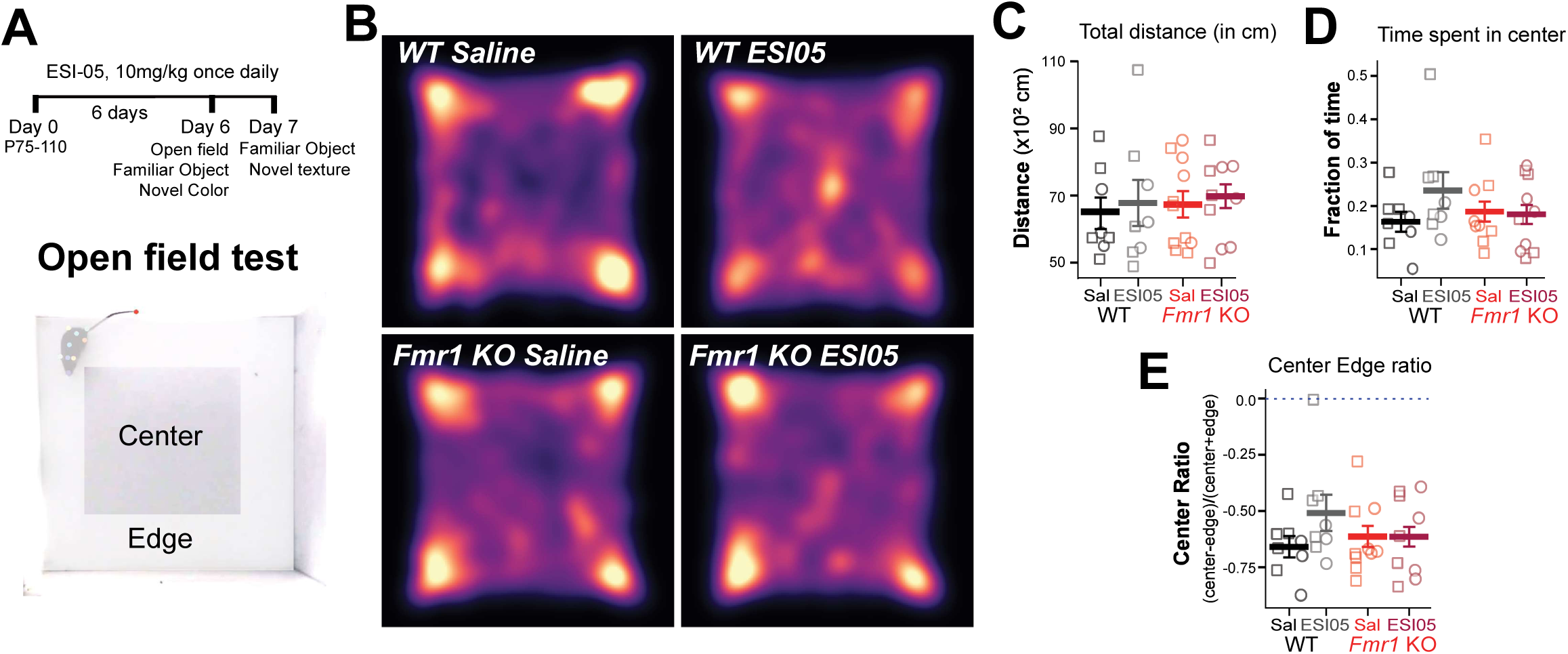
*Fmr1* KO and WT mice show similar behavior in no differences in open field test. **(a)** (Top) Experimental design and timeline for behavioral assessments. (Bottom) representative frame of movie recording open field behavior. **(B)** Mean heatmap of exploration during open field for WT and *Fmr1* KO mice treated with saline or ESI-05 (WT Saline=8, WT ESI-05=8, KO Saline=8, KO ESI-05=9). **(C)** Total distance travelled was similar across all groups. **(D)** The fraction of time spent in the center of the arena was similar across groups **(E)** The center edge ratio (a measure of how much time the animal spends in the edges of the arena) was similar across groups; the horizontal line at 0.0 indicates when mice would spend equal time in the center vs. the edges of the arena.

## METHODS

### Mouse lines

All experiments followed the U.S. National Institutes of Health guidelines for animal research, under an animal use protocol (ARC #2007-035) approved by the Chancellor’s Animal Research Committee and Office for Animal Research Oversight at the University of California, Los Angeles. All mice were housed in a vivarium with a 12/12 h light/dark cycle and experiments were performed during the light cycle. Animals were weaned from their dam at postnatal (P) day 21-22 and then group housed with up to five mice per cage. Mice of both sexes were used in this study (males are represented by a square and females by a circle). Because we use both sexes, we use *Fmr1* KO to describe both male (*Fmr1*^−/y^) and female (*Fmr1*^−/−^) knockout mice. The mouse lines used in this study were obtained from Jackson laboratory: *Pvalb*-Cre mice (JAX 008069), *Camk2a*-Cre (JAX 005359), Rpl22^HA/HA^ (JAX0110029). Mouse lines were crossed to WT (JAX line 000664) or *Fmr1* KO female mice (JAX 003025). For this study, the following mouse lines were generated: PV-Cre-*Rpl22*^HA/HA^-WT and PV-Cre-*Rpl22*^HA/HA^-*Fmr1*KO and *Camk2a*-Cre-*Rpl22*^HA/HA^-WT and *Camk2a*-Cre-Rpl22^HA/HA^-*Fmr1* KO. Transgenic lines were used as homozygotes for *Rpl22-HA*.

### RiboTag RNA extraction and sequencing

Adult (P70-100) *PV*-Cre-Rpl22^HA/HA^ and *Camk2a*-Cre-Rpl22^HA/HA^, WT or *Fmr1* KO mice were used for the Ribotag RNA-seq experiments. For each mRNA isolation experiment, mice were anaesthetized with isoflurane, followed by cardiac perfusion with ice cold choline ACSF containing 132 mM choline, 2.5 mM KCl, 1.25 mM NaH2PO4, 25 mM NaHCO3, 7 mM MgCl2, 0.5 mM CaCl2, and 8 mM D-glucose). Coronal sections (1.5 mm thick) containing S1 (A/P −0.5 to +1.82 mm, M/L 3.0 to 4.0 mm) and V1 (A/P +3.5 to 5.5 mm, M/L 1.5 to 3.0 mm) were collected using Stainless-Steel Brain Matrices (Zivic instruments) in chilled choline, from which S1 and V1 were dissected bilaterally. Samples were independently homogenized with a 2 mL Dounce homogenizer in 1 mL of supplemented homogenization buffer containing 1% w/v, HB-S; 1 mM DTT, protease inhibitors (1X), RNAsin (200 units/mL), cycloheximide (2 mg/l), and heparin (1 mg/mL). 100 μL of the homogenate was used for isolating bulk RNA from S1-V1. To isolate mRNA of HA-tagged *Pvalb*-Cre or *Camk2a*-Cre from non-HA tagged ribosomes, samples were incubated for 4 h with an anti-HA antibody (1:180, 5 μL in 900 μL of homogenate; 901513 Covance Anti-HA antibody), followed by overnight incubation with magnetic beads (Pierce Protein A/G Magnetic Beads 88802, 200 μL) at 4°C at 15 rpm centrifugation. The supernatant was washed 3 times with 800 μL of high-salt buffer (Tris 50 mM, KCl 300 mM, MgCl2 12 mM, 10% NP-40 (1%), DTT 1 mM, cycloheximide 100 μg/mL, dH_2_O), centrifuged at 15 rpm for 10 min at 4 °C. For RNA extraction, we used Zymo Research Direct-zol RNA Miniprep kit (R2051) for the homogenate (input), and the Zymo Research Direct-zol RNA Microprep kit (R2061) for the bead-antibody-protein sample (IP); RNA was eluted with DNAase/RNAse-free water. RNA Integrity Number (RIN) scores were used to evaluate the integrity of RNA through the ratio of 28S:18S ribosomal RNA (Agilent TapeStation 4200)^86^. We set a minimum RIN score of 7 for our samples. RNA-seq libraries were prepared using Ovation® RNA-Seq System V2. Libraries were multiplexed for paired end 100 bp sequencing on NovaSeq S2, with a read depth of 50 million reads on average. Demultiplexed reads were aligned using STAR aligner to the mouse genome (reference genome ID: mm10) and gene level quantification calculated by *Salmon*^87^. Outliers were calculated by measuring connectivity between samples, using the WCGNA package^46^ and removing samples >2 SD from the mean for Bulk and IP samples. All analysis was performed using R4.4.2. Differential expression analysis was performed using *voom* package was used to estimate the mean-variance relationship and adjust weights per animal basis. Differential gene analysis was performed using *limma*^35^ with FDR of 0.05. GO enrichment analysis and GSEA was performed using clusterProfiler 4.0.(enrichGO and fgsea) and SynGo for synaptic enrichment analysis^45,88^. For clusterProfiler analysis GO Terms with greater than 70% overlap of genes was combined into one category. The GO Term with the highest category was chosen as the name of the category. Pathways with adjusted p values >0.03 and non-brain related pathways were excluded with gene enrichment analysis. For comparisons with previously published RNAseq data, we compiled all DE genes from those studies^14,15,17,50,52,54^ and compared the compiled list with our DEGs. Statistical significance was tested using the Fisher exact test in R4.0.3.

### Network analysis

Network analysis was performed with the WGCNA package^46^ using signed networks. A soft-threshold power of 14 was used to achieve approximate scale-free topology (R2≅0.7). Networks were constructed using the *blockwiseModules* function. The network dendrogram was created using average linkage hierarchical clustering of the topological overlap dissimilarity matrix. Modules were defined as branches of the dendrogram using the hybrid dynamic tree-cutting method ^46^. Modules were summarized by their first principal component (ME, module eigengene), and modules with eigengene correlations of >0.9 were merged. Modules were defined using biweight midcorrelation (bicor), with a minimum module size of 100, deepsplit of 4, merge threshold of 0.1, and negative pamStage. Module differential expressions were determined using a linear model provided by the lmFit function in the Limma package^35^. Uncorrected *p* values are reported for module differential expression.

### Comparisons with other genes lists

*ASD risk genes:* A list of 1859 ASD risk genes were generated by combining candidate genes from the SFARI gene database (1203 genes, version 9 October 2024)^30^ and Geisinger Developmental Brain Disorder Gene Database (1233 genes, version: November 2024)^31^. We included genes that were candidates for both autism and ID from the Geisinger DBD and all genes irrespective of gene scores from the SFARI database.

*FMRP target* FMRP targets were identified from two previously published HITS-CLIP studies^10,12^. In the Maurin et al. dataset, only genes with total counts exceeding 50 across all examined brain regions were included.

*Brain enriched genes:* We evaluated mouse tissue enrichment on the CZ CELL×GENE Discover platform^89^ analyzing single cell data from >2 million cells, 31 tissue types from 43 research articles (as of February 2025). Embryonic tissue was excluded from the analysis. In the brain, average expression was determined as the mean expression for only neurons, whereas for all other tissues the expression was calculated for all cells within the tissue. We considered a gene as enriched if the tissue specificity index^90,91^ was greater than 0.7 and the brain had the highest expression.

*Excitatory and PV enriched genes:* For classifying genes as excitatory-enriched, we compared the trimmed mean expression per gene of all “CTX” subclasses with non-excitatory subtypes including Pvalb, Sst, Vip, Sncg, Oligo, Lamp5, Meis, Endothelial, Astrocytes, Pericyte, VLMC, CR and Pax6 as t-test with Bonferroni correction for multiple comparisons. Genes were considered as ‘enriched’ when the adjusted p-value was <0.03 and mean expression was 2-fold larger than non-cortical cell types and PV subtypes. For classifying genes as PV-enriched, a similar method was used except the subtypes compared included “CTX” and excluded *Pvalb* subtypes. There was a total of 255 genes for excitatory neurons and 92 genes for PV. Gene expression from non-cortical regions is not included.

*Comparison with histone modifier genes:* Gene lists were downloaded from MGI (Mouse genomic informatics) for biological processes related to histone modification activity (628 genes, GO:0140993) and compared with genes from the black module (Figure5A).

*Comparison with chromatin modifier genes:* Gene lists were downloaded from MGI (Mouse genomic informatics) for biological processes related to chromatin remodeling (1243 genes, GO:0006338). Inferred from electronic annotation (IEA) genes were excluded and only the following annotated terms chromatin remodeling’, ‘epigenetic programming of gene expression’, ‘heterochromatin formation’ and ‘epigenetic regulation of gene expression’ were included and compared with genes from *Camk2a*, *Pvalb* and bulk samples (Figure5B).

*Comparison with published proteomes*: We compared the DEGs in the *Camk2a* and *Pvalb* translatomes with previously published proteome datasets (Klemmer et al., 2011; Tang et al., 2015; He et al., 2019; Seo et al., 2023; Louros et al., 2023) and the FDR significance was set as per each study’s methodology (Table S2).

*Comparison with human FXS transcriptome data:* For cross-referencing with human data, we compared our DEGs to bulk RNA-seq data obtained from organoids, fetal tissue, and IPSCs derived from patients with FXS and isogenic controls^54^(Table S3). FDR significance was set as per that study’s methodology.

*Comparison with EPAC2 interactome*: EPAC2 interactome was generated from the BioGRID, String databases and from prior literature^92^.

### In situ hybridization

Mice were anesthetized with 5% isoflurane and transcardially perfused with ice cold phosphate buffer saline PBS (0.1 M) followed by ice cold 4% paraformaldehyde in PBS, post-fixed 12hrs at 4°C and then then transferred to sucrose gradients (10–15%, 20–22%, 30% sucrose in 1X PBS at 4 °C) and frozen in OCT. Coronal cryosections (20 µm) were cut on a Leica CM1850-3-1 Cryostat, mounted onto glass slides and stored at −80C. For RNAscope experiments, probes were obtained from ACD biotechne, Mm-*Rapgef4* #428851, Mm-*Pvalb*-C2 #421931-C2 and mM-*Slc17A7* # 416631-C3 and in situ hybridization performed using the RNAscope® Multiplex Fluorescent Reagent Kit v2 #323100. Staining was performed as per manufactures instructions with incubations done in Grace-Bio(#HBW2222FL) incubation chambers to prevent dehydration. Staining was done sequentially and imaged on an ApoTome2 microscope as 3 channels tiled zstack (Zen2 software, Zeiss; PAN-NEOFLUAR 10X objective, 0.3 NA). The section sampling fraction (ssf=1/6) for S1BF was determined within [A/P: −0.94 mm to −1.94 mm relative to Bregma]. For quantification, *Slc17A7* and *Pvalb* images denoised (python skimage medfilt2d (kernel size= 9), edge enhanced using skimage maximum filter followed by segmentation using *Cellpose* ^93^. *Rapgef4* puncta were quantified using the Find Maxima (prominence 25) function in Fiji. Overlap was quantified as overlapping of peak maxima with cell masks. To prevent overcounting of cells and puncta, overlaps across z were accounted for (as percentage overlap >0.8) and counts per cell calculated for the same cell across depth.

### Immunohistochemistry

Mice were anesthetized with 5% isoflurane and transcardially perfused with ice cold phosphate buffer saline PBS (0.1 M) followed by ice cold 4% paraformaldehyde in PBS and post-fixed overnight at 4°C. Coronal sections (40 μm) were cut on a vibratome (Leica VT1200S). For EPAC2 staining sections were incubated in a 1 mL PBS, 33 μL 30% H_2_O_2_, 3.3 μL 10% SDS solution at 60°C for 10 min. Sections were then blocked in 5% normal donkey serum (NDS) and Fetal Bovine Serum (FBS) for 30 min at RT. Sections were then incubated overnight at 4°C with the primary antibody diluted in PBS with 5% NDS and FBS and 0.005% Triton X-100. After rinsing in PBS for 5-10 min 3 times, sections were incubated for 1 h at RT with the corresponding Alexa Fluor-conjugated secondary antibody diluted in PBS (1:1000, Invitrogen). After rinsing in PBS for 5-10 min 3 times, sections were mounted onto Superfrost Plus glass slides (Vectashield Vibrance) with DAPI mounting medium and stored in the dark at RT. The following primary antibodies were used: mouse anti-parvalbumin (1:1,000, SAB4200545, Sigma Aldrich), Anti-EPAC2 (1:250, rabbitAb #43239 and mAb #4156, Cell Signaling), Anti-NEUN (1:1000, MA5-33103, Thermofisher), anti-HA (1:1000, mAb#901513, Covance), anti-GAD (1:1000, Gad6 DSHB), anti-VGLUT1 (1:1000, N28/9 DSHB). To evaluate overlap of PV, NEUN and HA, sections were imaged on an ApoTome2 microscope (Zen2 software, Zeiss; 20X objective, 0.8 NA), and analyzed with Fiji ^94^. The section sampling fraction (ssf=1/6) for S1BF was determined within [A/P: bregma −0.94 mm to −1.94 mm].

For EPAC2 immunostaining, images were acquired as 2-channel tiled (20 tiles) z-stacks on Zeiss LSM 700 confocal (Zeiss 100X PlanApo oil objective, 1.46NA, z-steps 0.5 μm) from pial surface to Layer 2/3 of S1 BF. For cell segmentation of NEUN cell bodies, images were denoised on Python skimage.medfilt2d (kernal size= 9) and cell segmentation was done on the NEUN channel for every plane using *Cellpose* cytotorch_1 model ^93^. Individual cells were tracked across Z’s by measuring overlap between masks with overlap >75% of the smaller mask was registered as the same cell. For EPAC2 puncta, images were processed in MATLAB (2023a) and ImageJ/FIJI using the MIJI plugin. Images were denoised (median filter,radius=2), background subtracted (rolling=5) and peaks identified (Find Maxima, prominence =100) using ImageJ/Fiji functions. Threshold was calculated as 1.5 SD of the maximal values and overlap between thresholded EPAC2 and cell bodies was quantified and calculated as a percentage for every cell and across Z planes. All samples were analyzed in an automated manner, blinded to genotypes and with similar settings to minimize experimenter bias.

### Synaptoneurosome (SN) preparation

We adapted a previously described protocol^95^. The mouse visual cortex at postnatal day (P) 14 was dissected, flash frozen, and stored at −80°C. Individual samples (approx. 1 mg) were homogenized in 700 µL SN lysis buffer (10 mM HEPES, pH 7.4, containing proteinase inhibitor) using a 2 mL Dounce tissue grinder (Kontes, Fisher K885300-0002) and applying 6-10 loose pestle (until disrupting all the tissue pieces) and 12 tight pestle strokes. A 150 µL aliquot of this total protein (TP) fraction was boiled in SN lysis buffer containing 10% SDS for 10 min and stored. The remaining fraction was centrifuged at 2000xg for 2 min at 4 °C to remove cellular and nuclear debris. The supernatant was loaded into a 1 mL syringe and filtered through three layers of pre-wetted 100μm pore Nylon filter (Millipore NY1H04700). The filtrate was directly loaded into 5 μm-pore centrifugal filters (Ultrafree-CL, Millipore UFC40SV25) and centrifuged at 5000xg for 15 min in a fixed angle rotor at 4 °C. The supernatant was carefully removed, and the loose pellet resuspended in boiling SN lysis buffer containing 2% SDS, boiled for 10 min and store at −80 °C (SN protein fraction). Protein concentration was determined with the BCA protein assay (Thermo Fisher 23227) using a bovine serum albumin standard curve.

### Co-immunoprecipitation

Flash frozen visual and somatosensory cortex from P25 (±1 d) WT mice was homogenized over ice using a 2 mL Dounce tissue grinder (Kontes, Fisher K885300-0002) in 400 µL of ice-cold lysis buffer containing 20 mM Tris (pH 7.4), 200 mM NaCl, 1% NP-40 and proteinase inhibitors. Protein was then extracted by incubating the homogenate on ice with occasional agitation for 30 min and then centrifuging at 15,000xg for 20 min. The pellet was resuspended in 200 µL lysis buffer and the extraction step was repeated. Both supernatants were pooled, and protein concentration was determined using the BCA method (Thermo Fisher 23227). Dynabeads protein G was used as suggested by the manufacturer (Thermo Fisher 10007D): Foreach sample 50 µL of magnetic beads were crosslinked to 2.5 µg of anti-PSD95 antibody (Antibodies Inc 75-028) or 2.5 µg of anti-SYT2 antibody (Abcam ab154035) with 230 µL of 5 mM BS3 (Thermo Fisher A39266). Samples were incubated with prepared beads for 16 h at 4 °C, 200µg of total pre-IP protein was used for PSD95 IP and 400 µg for the SYT2 IP, based on a pilot experiment showing low abundance of SYT2-associated protein. The supernatant containing the unbound proteins (post-IP) was stored and the beads washed. Finally, the bound protein (IP) was recovered in 10 µL elution buffer (50 mM Glycine pH 2.8) and used directly for Western Blot with the addition of denaturing loading buffer.

### Western blot

We followed a previously described protocol^95^. Protein samples were separated in Bolt 4-12% Bis-Tris Plus Gels (Thermo Fisher NW04125BOX) in Bolt MOPS SDS running buffer at 200 V for approximately 1 h. Except for SYT2 co-IP were samples were separated in NuPAGE 3-8% Tris-Acetate Protein Gels (Thermo Fisher EA03755) in Tris-Acetate-EDTA running buffer (Thermo Fisher LA0041) at 150 V for approximately 90 min. Separated proteins were transferred into a PVDF membrane using a Mini Blot Module (Thermo Fisher B1000) at 20 V for 1 hour except for Tris-Acetate gels transferred for 1:20h. Ponceau red staining was performed after transfer and recorded. Membranes were blocked in 0.1% Tween20 in Phosphate buffered saline (PBS) starting block (Thermo Fisher 37538) and incubated with primary antibodies diluted in the same buffer: anti-EPAC2 1:1,000 (Cell Signaling 43239S), anti-SYT2 1:1,000 (Abcam ab154035), anti-tubulin 1:10,000 (Sigma-Aldrich, T5326) and anti-PSD95 1:1,000 (Antibodies Inc 75-028). Signal was developed using ECL substrate (BioRad 170-5060) and recorded on a ChemiDocMP imager (BioRad). Densitometric analysis was performed with the open-source program FIJI and statistics performed on Prism10 (GraphPad).

### ESI-05 treatment in vivo

ESI-05, or 1,3,5-Trimethyl-2-[(4-methylphenyl)sulfonyl]-benzene, was purchased from Tocris (Cat no. 6320/25). Working stocks of ESI-05 were prepared by dissolving in molecular grade DMSO to a concentration of 20 mg/mL. Stocks were frozen and stored in −80°C. For experiments, this working stock solution was dissolved 1:1 in Tween 80 and then diluted 1:18 with sterile NaCl to get a final dose of 1 mg/mL. Adult mice were injected intraperitoneally (i.p.) with a dose of 10 mg/Kg body weight. Control animals were injected with vehicle only, consisting of DMSO in Tween 80 (1:1). Mice were injected once daily for 6-7 days while monitoring their weight and health. We did not observe any untoward effects of ESI-05 in mice similar to other studies using ESI05 ^79^. All experiments were carried out 1 hour after the treatment on the 6^th^ or 7^th^ day (for NORT texture experiments).

### Cranial windows and viral injections

Mice (P75-110) were anesthetized with isoflurane (5% induction, 1.5–2% maintenance via a nose cone, v/v) and their heads immobilized on a stereotaxic frame (Kopf) with ear bars. Carprofen (5 mg/kg, i.p., Zoetis) and dexamethasone (0.2 mg/kg, i.p., Vet One) were provided for pain relief and mitigation of edema, respectively, on the day of surgery and daily for the next 72LJhr. A 4.0 mm diameter craniotomy was performed over the left S1 cortex, under sterile conditions, using a pneumatic dental drill, taking care not to disturb the pia, as described previously ^96^. Stereotaxic viral injections of AAV1-Syn-GCaMP6s-WPRE-SV40 (Addgene 100843) were done using glass micropipettes (Sutter Instrument, 1.5 mm outer diameter, 0.86 mm inner diameter) through the open craniotomy. We used a Picospritzer (General Valve, 30 pulses of 6 ms, 30 psi) to pressure-inject 250-350nL of rAAV at 6-8 different locations over S1 cortex at a depth of 0.2 mm below the dura. The needle was left in place for 10 min to allow for diffusion and prevent backflow into the needle path. The craniotomy was then covered with a 5 mm glass coverslip and secured by cyanoacrylate glue and dental cement. A custom-made horseshoe-shaped titanium head bar (3.15 mm wide x 10 mm long) was attached to the skull (caudal to the window) with dental cement to secure the animal to the microscope stage. Within 1 h after surgery, when mice appeared fully recovered from the effects of anesthesia, they were returned to their home cage.

### Intrinsic signal imaging

For animals undergoing calcium imaging, we mapped the location of the barrel field within S1 using intrinsic signal imaging, 5-7 d after cranial window surgery, as described previously ^97,98^. The cortical surface was illuminated by green LEDs (535 nm) to visualize the superficial vasculature. The macroscope was then focused ∼300 μm below the cortical surface and red LEDs (630 nm) were used to record intrinsic signals, with frames collected at 30LJfps from 0.9 s before until 1.5 s after stimulation, using a 8 megapixel CCD camera (catalog #8051M-USB, Thorlabs), and custom routines written in MATLAB (version 2009a). Thirty trials separated by 20 s were conducted for each imaging session. The contralateral C2 whisker was gently attached with bone wax to a glass microelectrode coupled to a ceramic piezo-actuator (PI127-Physik Instrumente). Each stimulation trial consisted of a 100 Hz sawtooth stimulation lasting1.5 s. The response signal was divided by the average baseline signal, summed for all trials, with a threshold at a fraction (65%) of maximum response to delineate the cortical representation of stimulated whiskers and guide viral injections to S1BF using the same pipeline as described previously ^99^.

### In vivo 2-photon (2P) calcium imaging in head-fixed mice

In vivo calcium imaging was performed in awake mice. We used a commercial 2P microscope (DIY Bergamo, ThorLabs) equipped with galvo-resonant scanning mirrors, amplified non-cooled GaAsP photomultiplier tubes (Hamamatsu), a 16X objective (0.8NA, Olympus, ThorN16XLWD-PF), and ThorImage software. The microscope was coupled to a Chameleon Ultra II Ti:sapphire laser (Coherent) tuned to 930 nm, and the average power at the sample was kept <80 mW. We recorded both spontaneous activity and whisker-evoked activity in S1 guided by the location of the C2 map from the intrinsic signal imaging. Calcium imaging was performed at a framerate of 15.1 Hz. Mice were habituated to the microscope setup for 7 d prior to calcium imaging. This process lasted ∼5 d and involved a gradual progression from basic daily handling (5 min/d) until mice were comfortable with head fixation and body restraint in a plexiglass tube for periods up to 30 min. Mice were considered ready for imaging when they could remain still during sham whisker stimulation, in which the whisker stimulator is placed in front of the mouse but out of reach of its whiskers. The whisker stimulator consisted of a “comb” of von Frey Nylon filaments intercalated between whiskers (filaments were spaced 0.5 mm apart). This comb was coupled to a piezoactuator (controlled by MATLAB) that delivered repetitive deflections of the whiskers in the antero-posterior direction, with 1 s-long bouts of 20 Hz stimulation, and 3 s-long inter-bout intervals, for a total of 40 bouts but activity of the first 20 bouts were analyzed. To confirm that most whiskers on one side of the snout were being deflected consistently across different bouts, we used two cameras (FLIR BFS-U3-23S3M-C: Monocamera) at 30 and 90 fps to record both the position of the comb of Nylon filaments relative to the whiskers and body movement, pupil diameter and facial features of test mouse whiskers (Dobler et al., 2024) Blood vessels, visible as dark shadows, served as markers to consistently triangulate the same location over the 5-day period.

### Analysis of calcium imaging data

Calcium imaging data were analyzed using custom-written MATLAB routines (MATLAB version 2020a). Motion correction and ROI segmentation were done using Suite2p ^100^Automated ROI exclusion was performed using a custom classifier built using previous L2/3 data, followed by a manual refinement step to include or exclude ROIs that had been missed by the classifier. A ‘‘modified Z score’’ Z_F vector (an epoch was 8 consecutive frames wherein Z_F> 3 for every frame of that epoch) for each neuron was calculated as Z_F [F(t) mean(quietest period)]/SD(quietest period), where the quietest period is the 10 s period with the lowest variation (standard deviation) in dF/F, as described previously ^76^. Neurons were classified as active if they had at least one activity epoch with Z_F> 3 for 15 consecutive frames and only active neurons were used for subsequent analysis. To define whether an individual active neuron was stimulus responsive, we used a probabilistic bootstrapping (5,000 scrambles, threshold p<0.05, r^2 <0.2) method to identify neurons whose responses coincided with epochs of whisker stimulation as previously described ^76^. Adaptation of neuronal activity to repetitive whisker stimulation, was done similar to previous papers ^19,43,76^ where we calculated an adaptation index as follows: [(Z score during first five stimulations) - (Z score during last five stimulations)]/[(Z score during first five stimulations) + (Z score during last five stimulations). Negative vs. positive values for the adaptation index indicate facilitation or adaptation, respectively.

### Tactile defensiveness assay

We used a paradigm we previously established to assess tactile defensiveness to repetitive whisker stimulation in juvenile and adult mice ^19,43,76^. Briefly, titanium head bars were surgically implanted under anesthesia at P70-90 and then mice were habituated to head restraint and to running on an air-suspended 200 mm polystyrene ball. For habituation, mice were placed on the ball for 20 min/d for 5 consecutive days before testing. On the test day, each animal was first placed on the ball for a 3 min baseline period. Next, we performed a sham stimulation trial in which the whisker stimulator was visibly moving, but just out of the reach of the whiskers on the animal’s left side. The stimulator consisted of a long, narrow comb of five slightly flexible von Frey nylon filaments that were attached to a piezoelectric actuator (Physik Instrumente). During the stimulation trial, the filaments were intercalated between the whiskers. Bundling whiskers onto a glass capillary was not feasible for these awake experiments because the mouse could have damaged the capillary or unbundled some of its whiskers with its forepaw. The stimulation protocol consisted of a 10 s baseline followed by 40 sequential bouts of whisker deflections along the anterior–posterior direction (1 s long at 20 Hz), with a 3 s inter-bout interval, and ending with another 10 s post-stimulation baseline. We used a wide angled webcam (EMEET C960,1080p, 30 fps) camera to record the responses of the animal. A custom-written semiautomated video analysis was implemented in MATLAB to score defensive behaviors (grabbing the stimulator) and adaptive healthy behaviors (grooming), in each 1s increment of the videos during the 20 stimulations.

### Open field and novel object recognition tasks (NORT)

All behavior experiments were carried out in a custom-built white square HDPE box (40 x 40 cm) under dim lighting (18 lumen). Mice (7-10 weeks) were recorded using a wide angled webcam (EMEET C960,1080p, 30 fps) placed above the box. For NORT experiments the location of the novel object was randomly assigned. Movies were then converted to *.mp4, trimmed to 10 min using custom Python scripts and the *ffmpeg* library. Movies were then individually cropped and downsized (390 X 390), rotated to align the movies (so the novel object is always in the same location for analysis), and brightness/contrast-adjusted (to aid in *DeepLabCut* training) using the *Shotcut* software. Videos were then trained on individual networks using *DeepLabCut* ^82,83^ (resnet 50, shuffle 1, iterations 30000). Markers placed on the nose, head, ears, neck, upper back, lower back, forelimbs, hindlimb, base and tip of tail were tracked simultaneously. Additionally, features of the behavioral arena, such as corners and edges of objects, were also tracked by a separate trained network. The position of each mouse was calculated as the average location of its head for every second of recording (a mean of 30 frames). The location of the mouse was calculated relative to the centroid of the box, with the ‘central region’ of the arena defined as a square area measuring 16 x 16 cm around the centroid and corners defined as smaller squares (5 x 5 cm) at each corner of the arena. The center of the chamber was calculated as a centroid of the four corners of the chamber and care was taken to position the webcam such that all movies were centered around the centroids. We then calculated the proportion of time each mouse spent in the central or corner regions. Experiments were carried out across two days, with open field, object habituation, and novel object recognition task-visual being tested on Day 1 and object habituation and novel object recognition task-texture being tested on Day 2. Mice returned to their home cage after each task and tested on the next task after a break of 15 min. The chamber was wiped with 70% ethanol and water before testing the next mouse. Before every experiment all objects were rinsed with water, ethanol, and water again before being thoroughly dried.

#### Open Field

Naïve mice (7 to 10 weeks) were gently placed in the center of the box and allowed to freely explore for 10 min. We calculated the total distance traveled and the time spent in the central region during the 10 min video.

#### Object habituation

Two black acrylic cubes (5 x 5 x 5 cm), each with one face wrapped in red tape oriented toward the camera, were placed equidistant from each other and the arena edges. Mice, previously familiarized with the objects, were placed in the center and allowed to explore freely for 10 minutes, with tracking enabled by DeepLabCut. The red tape on the object was to provide sufficient contrast for DeepLabCut to track while the mouse was on top of the cubes. In Figure 8, the surface with the red tape is covered in black for illustration purposes.

#### NORT-visual

We replaced one of the familiar red cubes from the previous object habituation task with a novel yellow plastic cube of equal dimensions. The location of the novel object was randomized for each experiment. Mice were placed gently in the center of the chamber and allowed to freely explore the box for 10 minutes.

#### NORT-tactile

On Day 2, following object familiarization, one of the familiar cubes was replaced with a novel cube of the same color and dimensions but wrapped in Velcro. The location of the novel object was randomized for each experiment. Mice were placed gently in the center of the chamber and allowed to freely explore the box for 10 min.

### Software and statistical analysis

We used MATLAB (2023a), ImageJ/FIJI, and MIJ for data processing and visualization. We used R (4.2.3) and GraphPad Prism 9/10 for statistical analyses, as indicated in the text. \*\**p*-value<0.01, \*\*\**p*-value<0.001, \*\*\*\**p*-value<0.0001, unless stated otherwise. Central tendencies were reported as mean of mice ± standard error for **Figures 3(c,f), 7(d-e, g-i), 8(c-d, g-h), 8f, 9(c-e),** median±1.5*intra-quartile range of mice for **Figures 1d, 6(d,f,h), 1c**. Tests for normality were tested for all data using Shapiro-Wilk’s method. For **Figure 7(g-i)**, data was log-normalized before analysis to satisfy the prerequisite assumptions of normality. All statistical tests are reported in Figure legends, and corresponding adjusted p-values are reported in Figures. All enrichments of gene sets were performed using a two-sided Fisher’s exact test with 95% confidence calculated using the R fisher function.

## Data and code availability

All data reported in this paper will be shared by the lead contact upon request. The original code has been deposited at github.com/porteralab as listed in the key resources table. Any additional information required to reanalyze the data reported in this paper is available from the lead contact upon request.

## Acknowledgements

The authors thank Irma Tello García, Toshihiro Nomura, John N. Armstrong, Li Yuan, Diana Mitchell, Sabrina Tazerart, and William Zeiger for technical assistance and/or suggestions throughout the project, as well as Anis Contractor, Anna Dunaevsky, and Padmashri Raghunathan for feedback on the manuscript. This work was supported by the National Institute of Neurological Disorders and Stroke (R01NS117597 to C.P.-C.), the National Institute of Child Health and Development (R01HD108370 and R01HD054453 to C.P.-C.; P50HD103557-05 to M.J.G.), the National Institute of Mental Health (R01MH121521 and R01MH137578 to M.J.G.), the Department of Defense (DOD 13196175 to C.P.-C.), the National Science Foundation (Award #2225624 to M.J.G.), the Canadian Institutes of Health Research (CIHR grant MOP-377520 to R.A.), an internal grant from the Center for Autism Research and Treatment and the Clinical and Translational Science Institute at UCLA (10POR2020 to C.P.-C.), a Marion Bowen Neurobiology Postdoctoral Fellowship (to A.S.), and the FRAXA foundation (post-doctoral fellowship to N.K.). We are grateful to the TGCB Sequencing core and the BSCRC confocal core, and to Graziella Di Christo for the SYT2 antibody.

## Author contributions

Conceptualization: A.S., M.J.-G. and C.P.-C. Methodology: A.S., J.E.B., and A.T.T. for RNA-seq experiments and analysis; R.A. and S.M.R for biochemical experiments; A.S. and C.S.L. for calcium imaging; A.S. and N.K. for behavioral experiments; A.S. and L.T.W. for histology, immunohistochemistry and in situ hybridization; A.S., M.J.-G. and C.P.-C. wrote the manuscript with input from other authors.

